# Strain stiffening of Ndc80 complexes attached to microtubule plus ends

**DOI:** 10.1101/2022.03.15.484438

**Authors:** Felix Schwietert, Vladimir A. Volkov, Pim J. Huis in ’t Veld, Marileen Dogterom, Andrea Musacchio, Jan Kierfeld

## Abstract

In the mitotic spindle, microtubules attach to chromosomes via kinetochores. The microtubule-binding Ndc80 complex is an integral part of kinetochores, and is essential for kinetochores to attach to microtubules and to transmit forces from dynamic microtubule ends to the chromosomes. The Ndc80 complex has a rod-like appearance with globular domains at its ends that are separated by a long coiled coil. Its mechanical properties are considered important for the dynamic interaction between kinetochores and microtubules. Here, we present a novel method that allows us to time-trace the effective stiffness of Ndc80 complexes following shortening microtubule ends against applied force in optical trap experiments. Applying this method to wild type Ndc80 and three variants (CH-domains mutated or Hec1-tail unphosphorylated, phosphorylated, or truncated), we reveal that each variant exhibits strain stiffening, i.e., the effective stiffness increases under tension that is built up by a depolymerizing microtubule. The strain stiffening relation is roughly linear and independent of the state of the microtubule. We introduce structure-based models, which show that the strain stiffening can be traced back to the specific architecture of the Ndc80 complex with a characteristic flexible kink, to thermal fluctuations of the microtubule, and to the bending elasticity of flaring protofilaments, which exert force to move the Ndc80 complexes. Our model accounts for changes in the amount of load-bearing attachments at various force levels and reproduces the roughly linear strain stiffening behavior, highlighting the importance of force-dependent binding affinity.

**SIGNIFICANCE:** By time-tracing the stiffness of microtubule end-tracking Ndc80 complexes in optical trap experiments, we detect strain stiffening, and, thereby, provide new insights into the elastic properties of the Ndc80 complex. The strain stiffening is robust against mutations in the Ndc80 complex. We relate strain stiffening to the structure of the Ndc80 complex by means of a simple polymer model, to thermal fluctuations of the microtubule, and to the flexibility of force-generating flaring protofilaments at the tip of the microtubule. Since Ndc80 complexes play a major role for transmitting force from microtubule ends to the kinetochore, their elastic properties are of great interest for a deeper understanding of chromosome dynamics in the mitotic spindle.

## INTRODUCTION

In the mitotic spindle, microtubules (MTs) attach to chromosomes via macromolecular structures called kinetochores (1). Kinetochores bind to MTs via attachments created by Ndc80 complexes (2). The Ndc80-mediated attachments remain intact while dynamic MTs alternate between growth and shrinkage, and transmit depolymerization forces to the kinetochore during the segregation of chromosomes. The exact mechanisms underlying binding and force transmission are not completely understood but are expected to reflect the molecular structure and resulting elastic properties of the Ndc80 complex.

The Ndc80 complex is a 4-subunit rod-like coiled-coil with a characteristic flexible kink at approximately one third of its length (3, 4). MT binding is performed by the globular calponin homology (CH) domains near the N-terminal ends of the Ndc80 and Nuf2 subunits, and an unstructured N-terminal tail. In vitro experiments reveal that a single Ndc80 complex is not able to track dynamic MT ends (5–7), whereas tracking activity is attained by multimerization (5, 7) and in presence of Dam1 or Ska complexes (6, 8, 9). Moreover, an unstructured positively charged Ndc80 tail that precedes the CH domain is crucial for MT tip tracking (10), and phosphorylation of the tail or introduction of negative charges reduces the binding affinity of the Ndc80 complex to the MT lattice (2, 11–14).

Early theoretical models of spindle-kinetochore dynamics envisioned a static Hill sleeve with rigid linkers (15, 16) or motor proteins mediating MT–kinetochore attachment and movement (17). More recent studies included (visco)elastic MT– kinetochore linkers with a constant stiffness, which had to be estimated (18–21). One such model identified the linker stiffness as a critical parameter for the occurrence of chromosome oscillations during metaphase (22). With the identification of the Ndc80 complex as the crucial linker, understanding its elastic properties has now become crucially important to dissect how MT depolymerization forces are transmitted to the kinetochore to generate movement. The elastic properties of the Ndc80 complex in its MT-bound state, in particular its stiffness under tension, are the main subject of this work.

Measurements of stiffness can confirm structural models if they are in agreement with elastic models derived from the available structural information on the Ndc80 complex. The elastic properties of the Ndc80 complex were studied experimentally by optical trapping methods introduced in Ref. 7. Here, we re-analyzed them using novel theoretical analysis and modeling approaches. In those experiments, the stiffness of Ndc80 complexes was found to increase under tension (7). Whereas in those experiments the stiffness was only determined while the Ndc80 complexes were attached to a stalled MT, here, we re-evaluate the same data with a novel method that allows us to time-trace the stiffness, i.e., to determine the stiffness of Ndc80 complexes while it tracks a polymerizing or depolymerizing MT against the opposing load from the optical trap. By splitting the force traces into fixed time intervals, and analyzing stiffness at each of them, we achieve several advantages: (1) we generate more data over a wider force range; (2) we demonstrate stiffening during force production and not only compare stalled MTs to free ones; (3) we alleviate the concern that different levels of strain stiffening result from differences in MT ends or beads. Time-tracing also allows us to establish that strain stiffening does not depend on the the state of the MT end: we observe positive correlation of stiffness with force when the MT stalls, grows, or shrinks. In addition to wild type Ndc80, we also analyze data from MT end-tracking experiments with Ndc80 complexes whose tails are phosphorylated or truncated (10), or harboring CH domains that are mutated to greatly reduce MT binding. We observe strain stiffening for each of those Ndc80 mutations and draw conclusions on the role of the tail domain in MT binding.

In order to rationalize the experimental findings on strain stiffening of the Ndc80 links, we introduce a simple polymer model for the Ndc80 complex. Describing Ndc80 as an ideal chain with two bonds of different lengths, we show that the strain stiffening is a direct consequence of the characteristic Ndc80 structure with its two stiff rods that are connected flexibly. To match the measured stiffnesses it is necessary to include the contributions from MT fluctuations and from protofilament (PF) bending to the total effective stiffness. Finally, to explain the shape of the experimental stiffness–force relations, we also introduce a positive correlation of the number of MT end-attached Ndc80 complexes with MT-generated force.

## MATERIALS AND METHODS

### Experimental methods

#### Cloning and purification

The Ndc80 CH-mutant contains Hec1 K89A, Hec1 K166A, Nuf2 K41A, and Nuf2 K115A: four mutations that reduce the binding of Ndc80 to MTs (23). Mutations were introduced into pLIB-Hec1 and pLIB-Nuf2 vectors using site-directed mutagenesis and Gibson assembly. Expression cassettes containing mutated Hec1, mutated Nuf2, Spc25-Sortase-Hexahistide and Spc24-Spy were combined on a pBIG1 plasmid and integrated into a single baculovirus for expression in insect cells (24). The Ndc80 CH-mutant was purified and linked to T_1_S_3_ modules (25) as previously described in detail for wild type, tailless, and phosphorylated Ndc80 variants (7, 10).

#### Optical trap experiments

Optical trap experiments with dynamic MTs and Ndc80 were performed as described previously (7, 10). In brief, 1 µm silica beads were coated with poly-L-lysine-grafted polyethyleneglycole (PLL-PEG) as described previously (7). 1–10 % of the PLL-PEG carried a biotin, and was subsequently saturated with T_1_S_3_[Ndc80]_3_ modules. Ndc80-TMR fluorescence on the bead was measured to determine the coating density, and bead preparations containing on average 100–1000 Ndc80 copies per bead were used for optical trapping.

The microscopic flow-chambers were assembled using glass slides and coverslips treated with 2 % dichlorodimethylsilane. The chambers were incubated with 0.2 µM of anti-digoxigenin IgG (Roche). Non-specific binding of molecules to coverslips was prevented by an incubation with 1 % Pluronic F-127. Then, GMPCPP-stabilized, digoxigenin-labeled MT seeds were attached to the coverslips. Finally, T_1_S_3_[Ndc80]_3_-coated beads were introduced in a reaction mix containing 10–12 µM tubulin, 1 mM GTP, 1 mg/ml *κ*-casein, 4 mM DTT, 0.2 mg/ml catalase, 0.4 mg/ml glucose oxidase and 20 mM glucose.

Optical trapping experiments were performed as described previously (7). Before each experiment, a free bead was attached to a MT near its growing end and held in an optical trap with a stiffness of 0.02–0.04 pN nm^*-*1^. Position of the bead was continuously recorded at 10 kHz using a quadrant detector, and differential interference contrast (DIC) images were simultaneously recorded to monitor positions of the bead and the dynamics of MTs. The bead was held in a trap until the MT depolymerized, which resulted in two alternative outcomes: the bead–MT connection broke, or the bead rescued the MT shortening, and the MT started regrowing. In case of a rescue, recordings were stopped arbitrarily after 30 or more minutes by increasing the trap stiffness to 0.2–0.4 pN nm^*-*1^ and rupturing the bead–MT connection. Quadrant detector readings of the bead position were later turned so that one of the axes corresponded to bead displacement along MT to generate time traces *x*(*t*) of the bead position.

### Time-tracing the stiffness

As long as the mean bead position is stationary, e.g. during MT stall, the effective stiffness can be easily calculated from the variance of the bead position *x* by *k*_B_*T*/Var(*x*). We now present a method that enables us to monitor the stiffness over time, i.e., also during polymerization and depolymerization phases of the MT, and to observe strain stiffening of the Ndc80 complex in the course of a single experiment. We divide the signal into several intervals *k* of a certain length Δ*t* for each of which we will determine a stiffness as sketched in Fig. 1. For that purpose, we do not use the variance over that interval but the mean squared distance from a quasi-equilibrium bead position which may slowly vary over time. We estimate the time course of the quasi-equilibrium bead position with a linear fit *f*_*k*_(*t*) to the entire time trace *x*_*k*_(*t*) in an interval *k*. Then, the effective stiffness during the interval *k* is given by the inverse of the mean squared distance

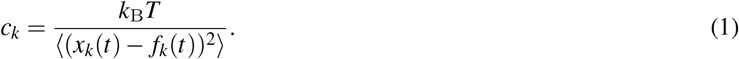

**Figure 1:**
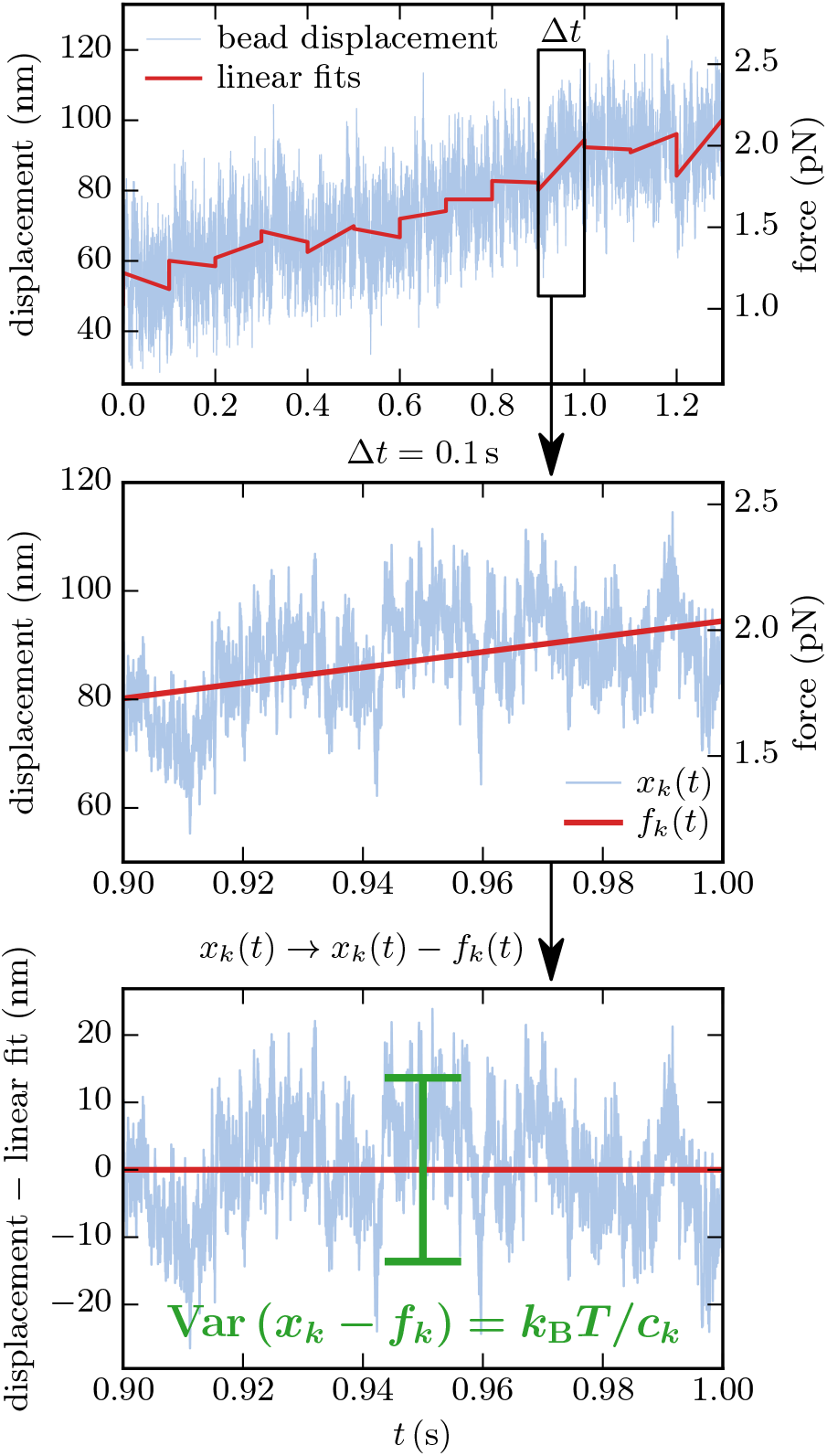
Stiffness time-tracing. The signal is divided in intervals of 0.1 s, which gives 1000 data points per interval at a sampling rate of 10 kHz. For each interval, we estimate the evolution of the equilibrium bead position with a linear fit *f*_*k*_(*t*). The stiffness of an interval is then given by Eq. 1, where the variance equals the mean squared distance of the bead position *x*_*k*_(*t*) from the estimated equilibrium *f*_*k*_(*t*).

Finally, we can assign a force *F*_*k*_ on the Ndc80 link to each interval, which can be calculated from the trap stiffness and the mean bead position during that interval, *F*_*k*_ = *c*_trap_ *x*_*k*_).

The time interval Δ*t* should be sufficiently small to describe a time-resolved stiffness but, at the same time, should cover enough independent displacement measurements for a trustful variance determination. Typical autocorrelation times are *γ/c* ∼ 0.1–1 ms, where *γ* is the friction coefficient and *c* the stiffness. For our experiments, Δ*t* = 0.1 s has proven to be a good choice, which results in 100-1000 independent measurements within each interval.

### Theoretical Model

Before modeling the outcomes of the stiffness measurements, we have to identify the elastic elements that contribute to the total stiffness as sketched in Fig. 2B. The stiffness of the optical trap will always be present in the measurements. Its value is determined during the calibration of the optical trap after the bead detached from the MT.

**Figure 2:**
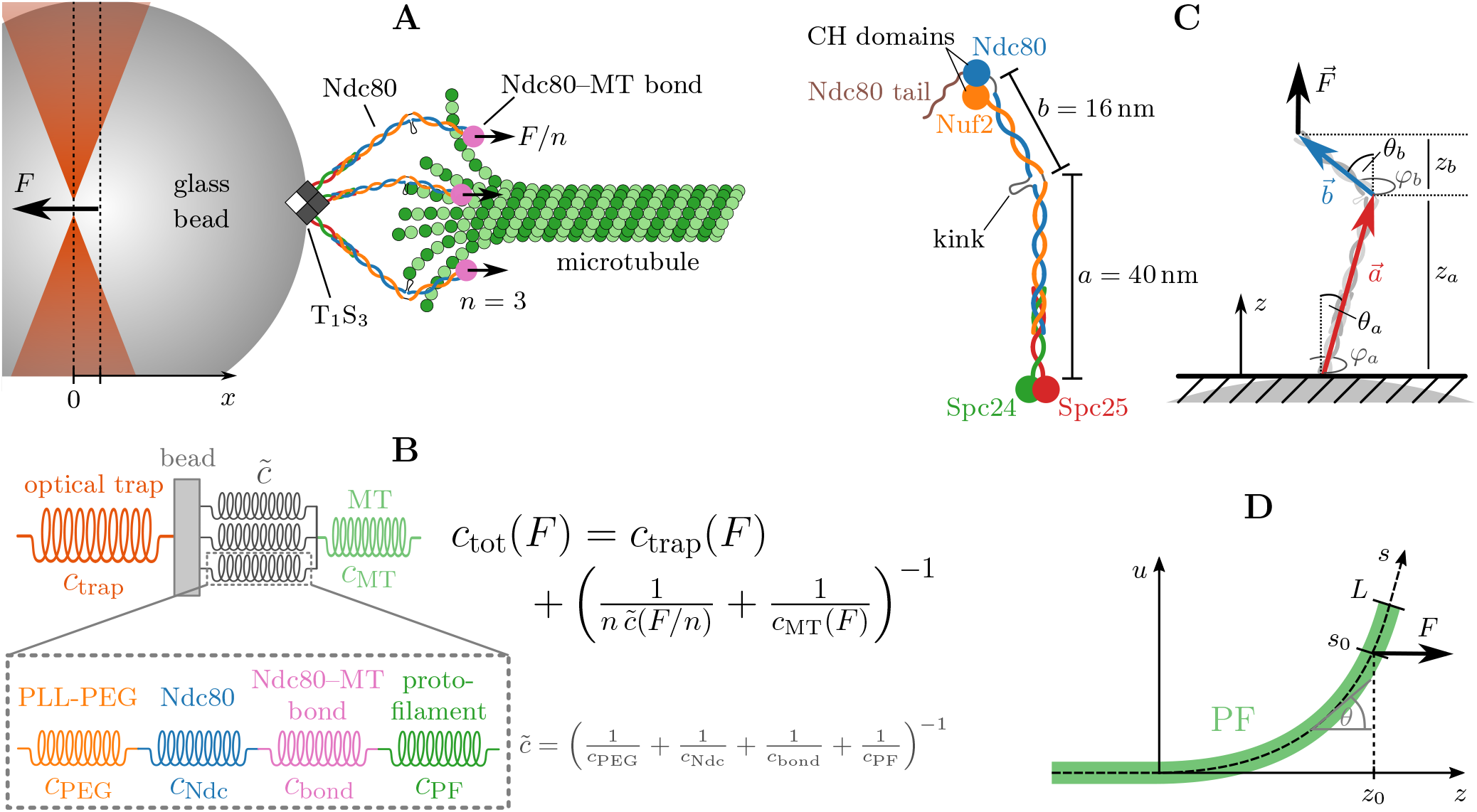
(A) Sketch of the experiment (not true to scale) with *n* = 3 attached Ndc80 complexes. We assume that each Ndc80 complex is exposed to the same force *F/n*. (B) Spring model. Between the bead and the MT, there are four potentially elastic objects, which are aligned in series: the PLL-PEG, the Ndc80 complex, the Ndc80–MT bond and the curved PF. When *n* Ndc80 complexes are attached to the MT, *n* of those serial combinations act as parallel springs. (C) Ndc80 complex as FJC. The Ndc80 complex is a tetramer with a total length of 56 nm and a flexible kink at 40 nm. The long arm *a* is bound to the glass bead, the short arm *b* to the MT. We model the Ndc80 complex as a FJC with two bonds 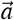 and 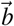, which have the different lengths *a* and *b*. (D) Model for PF bending. The *z*-axis marks the direction along the MT, *s* is the position along the curved PF. The Ndc80 complex is attached at *s* = *s*_0_ and applies the force *F* in *z*-direction.

Further potentially elastic elements connect the bead and the MT seed that is fixed on the coverslip. On the one hand, this is the MT, which can exhibit both a mechanical (stretching or bending) and an entropic stiffness (from thermal fluctuations). On the other hand, the link between the MT and the bead has elastic properties. We attribute the elasticity of the link to its four constituents: first, the Ndc80 complex can act as an entropic spring with stiffness *c*_Ndc_ due to its flexible kink; secondly, stretching the flaring ends of the PFs at the MT tip out of their preferred curvature may produce an effective stiffness *c*_PF_ for pulling in the axial direction of the MT; thirdly, also the bond between the Ndc80 trimer and the MT may exhibit some effective elasticity *c*_bond_, e.g. as a consequence of unbinding and rebinding of individual Ndc80 units; and finally, the PLL-PEG that connects the Ndc80 complexes with the bead is a flexible polymeric linker with an entropic stiffness *c*_PEG_.

Each of these four stiffnesses may depend on the applied force and exhibit strain stiffening itself. Since the four elements are aligned in series, their inverse stiffnesses sum up to the inverse linker stiffness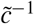:

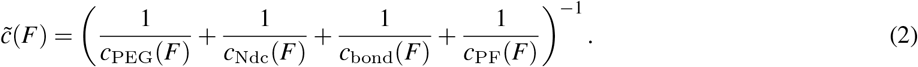

It is important to note that this reduces the total stiffness and that the softest elastic element dominates the total stiffness.

It is possible that multiple Ndc80 complexes are attached parallelly to the MT. For the sake of simplicity, we will always assume that each PF can only attach one Ndc80 complex and that the parallel PFs and Ndc80 complexes have the same elongation, respectively. Then, the force *F* is shared equally between the parallel Ndc80–PF units with stiffness 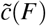, so that the resultant stiffness of *n* parallelly attached Ndc80 complexes can be written as

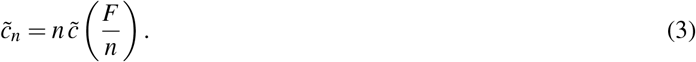

Although parallel stiffnesses add up, the shared force can give rise to an overall reduction 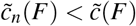 of the cooperative stiffness if the individual stiffness exhibits strain stiffening *c*(*F*) ∝ *F*^*m*^ with an exponent *m* > 1. We will find below that such strain stiffening behavior is realized both for Ndc80 and PF stiffness.

The *n* parallel linkers 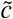 are in series with a single MT with stiffness *c*_MT_, and the optical trap, whose force is applied on the bead from the opposite side, acts as a parallel spring, see Figs. 2AB. This results in a total stiffness

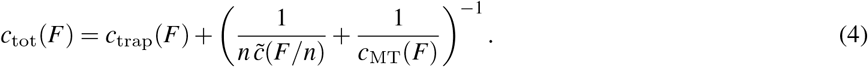

We note that the force dependence of the trap stiffness is negligible in the examined force ranges so that we can assume *c*_trap_(*F*) = const in the following.

#### MT stiffness

In the experiments, typical MT lengths *L* (i.e., the distances between the fixed MT seed and the MT tip linked to the bead) lay in a range between 3 and 10 nm. Given the values of 0.1 to 2 GPa for the Young’s modulus *Y* of a MT (26), the stiffness for MT stretching can be estimated as *YA/L* = 5 to 330 pN nm^*-*1^, which is stiff enough to be ignored in the following. Since the MT tip is lifted from the coverslip and therefore slightly bent when it is bound to the bead, a horizontal force might not only stretch but also unbend the MT. The mechanical stiffness from this unbending can also be neglected as we show in the Supporting Material.

Apart from purely mechanical elasticity, the MT can also exhibit an entropic stiffness, which follows from a description as a semiflexible polymer with thermally excited bending fluctuations. Then, for large forces (*F* ≫ *k*_B_*T/L*_p_ ∼ 10^*-*6^ pN), the relation between the applied force *F* and the mean extension *z* in force direction is given by (27, 28)

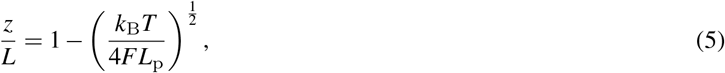

with the persistence length *L*_p_, which depends on the MT length (29): 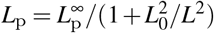 with 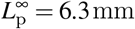 and *L*_0_ = 21 µm. The entropic MT stiffness *c*_MT_ can be deduced from the derivative of *z* with respect to the force:

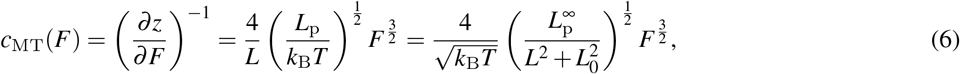

where the length-dependent persistence length was used in the last step.

### Bond stiffness

The bond between individual Ndc80 complexes and the MT is established by the globular regions at the N-terminal of the complex (3, 23, 30) via calponin-homology (CH) domains (31). Also the N-terminal tail of the Ndc80 subunit could be involved in MT binding (1, 32). The effective stiffness of the Ndc80–MT bond is hard to model because the exact binding mechanism to the MT is still elusive. Because the Ndc80 CH domain and the N-terminal tail are relevant for binding to the MT, investigation of the stiffness changes for mutants Ndc80^Δ80^ and Ndc80^CHmut^ as compared to the wild type Ndc80^wt^ will allow us to address this issue in the Discussion section. Regarding the linker stiffness 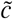, we will first concentrate on the contributions of the Ndc80 complex itself, the PF and the PLL-PEG in the following.

#### Ndc80 stiffness

In order to model the stiffness of the Ndc80 complex under force, we use a simple polymer model that is based on the known structure (3, 4): The Ndc80 complex is a tetramer with a total length of 56 nm and consists of two stiff arms with lengths *a* = 40 nm and *b* = 16 nm, see Fig. 2C. In the cell, the long arm is bound to the kinetochore, in the experiment it is bound to the glass bead, while the short arm can attach to a MT. We describe the conformation of the two arms by two vectors 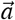 and 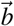 in polar coordinates as shown in Fig. 2C:

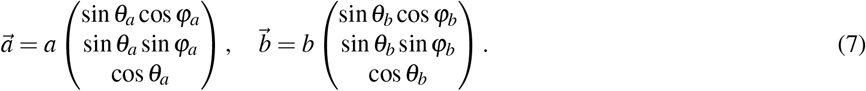

The impenetrable glass bead restricts these two vectors in the experiment. Since the diameter of the glass bead (1 µm) is much larger than the Ndc80 complex we approximate the bead as a plane surface that confines the Ndc80 complex to the upper half space, so that *z*_*a*_ > 0 and *z* = *z*_*a*_ + *z*_*b*_ > 0, see Fig. 2C. Experiments have shown that the two arms are connected flexibly within an angular range between 0° (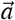 and 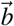 in line) and 120° (maximally kinked) (4). Since we want to model the Ndc80 complex while it is confined by the glass bead and under the influence of an external stretching force that is applied in *z*-direction and favors small angles, we can neglect the constraint to angles below 120°. This allows us to perform an explicit analytical calculation, as we reduce the Ndc80 complex to a purely entropic freely jointed chain (FJC) with two bonds of different lengths. The only energy is the stretching energy from a constant external force 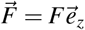 in positive *z*-direction,

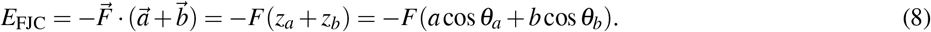

Respecting the confinement to the upper half-space we obtain the canonical partition function (*β ≡* 1/k_B_*T*):

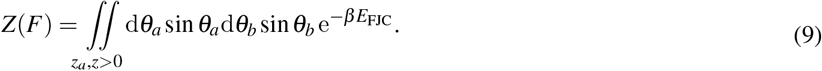

Both the extension–force relation *z*(*F*) and the stiffness–force relation *c*_Ndc_(*F*) in the presence of thermal fluctuations can be obtained from derivatives of the partition function with respect to the force:

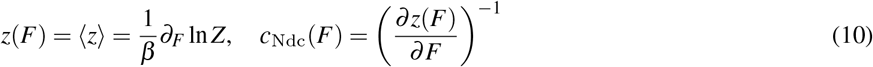

The explicit result for the stiffness of the Ndc80 complex is

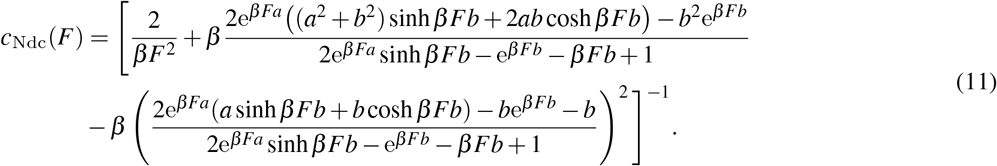

In the limit of strong forces (*F ≫* 1/βb = 0.256 pN), this simplifies to

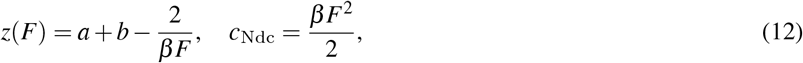

which is a general result for arbitrary FJC polymer models under force, where the relative extension of a FJC of total length *L* and bond length *b* saturates according to 1 *-z*(*F*)/L ∝ *k*_*B*_*T/bF* (28, 33).

#### Protofilament stiffness

We assign a stiffness to the PF by modeling it as a beam with a preferred curvature *ϕ* and persistence length *L*_p_. As described in Fig. 2D, the PF is straight and bound by lateral interactions with neighboring PFs for *z* < 0. For *z* > 0, we assume that there are no interactions to neighboring PFs so that the PF can bend freely. The PF conformation is described by the local bending angle *θ* (*s*) at the position *s* along the PF. The PF curvature is obtained by the derivative 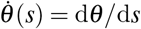. In the absence of an external force, the PF has a preferred (or spontaneous) constant curvature *ϕ*. Let the Ndc80 complex attach at the position *s* = *s*_0_ and apply a force *F* in *z*-direction. Then, the total energy is the sum of bending and stretching energies,

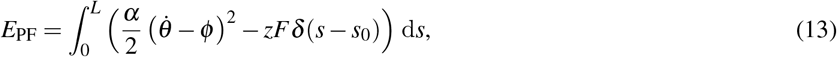

with the bending stiffness *α* = *k*_B_*TL*_p_ and the total length *L* of the free PF; the *δ* -function describes a point force applied at *s* = *s*_0_. The PF is dominated by the interplay of bending and stretching energy such that we can neglect thermal fluctuations and obtain the PF stiffness from the configuration minimizing the total energy *E*_PF_. Since the force *F* only affects the shape of the PF for *s* < *s*_0_ while the curvature at *s* > *s*_0_ stays *ϕ*, we have to minimize the PF energy with boundary conditions *θ* (0) = 0 and 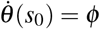. The energy minimizing PF configuration satisfies the Euler-Lagrange equation 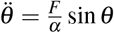, which we solve numerically with a shooting method in order to fulfill the boundary conditions. The effective deflection is given by the position of the point of force application on the *z*-axis *z*_0_ = *z*(*s*_0_). Finally, the effective stretching stiffness of the PF can be determined from the derivative of the deflection:

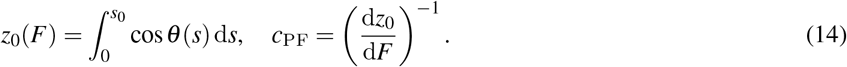

In the limit of strong forces 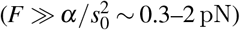, the PF is stretched, i.e., *θ* (*s*) ≪ 1, and we can approximate the differential equation by 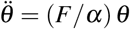, which can be solved analytically:

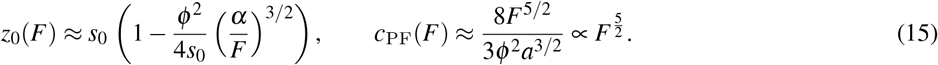

We note the pronounced strain stiffening with *c*_PF_(*F*) ∝ *F*^5/2^, which is a consequence of bending rigidity together with spontaneous curvature. The model depends on three parameters: The persistence length *L*_p_, the preferred curvature *ϕ*, and the point of force application *s*_0_. Following the results of in vitro experiments (34), we use *L*_p_ = 200 nm and *ϕ* = 20° per dimer throughout the paper. We assume that the Ndc80 complex can attach to the curved free part of the PF at a point *s*_0_ that has to be smaller than the length *L* of the curved part of the PF. This assumption will be revisited in the Discussion section. McIntosh *et al*. (34) measured lengths *L* in the range of 10 to 80 nm for depolymerizing MTs in vitro. In this paper, we will use estimates *s*_0_ = 20 nm and *s*_0_ = 50 nm.

#### PEG stiffness

We estimate the PEG stiffness on the basis of the extended FJC model for the elasticity of PEG provided by Oesterhelt *et al*. (35). Since PEG that is dissolved in water undergoes a conformational transition from a helical to a planar structure when tension is applied, the model includes two distinct bond lengths, one for each of the conformations. Assuming a Boltzmann distributed ratio of helical and planar monomers, the authors obtain an effective bond length that depends on the applied force. We derive the PEG stiffness *c*_PEG_ from the extension–force relation *L*(*F*) that is given in Eq. 2 in Ref. 35:

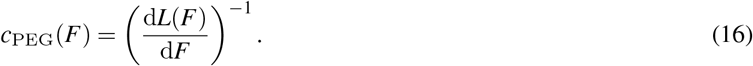

For the PEG-2000 used in our experiments, we evaluate the equations with a number of *N*_S_ = 45 segments while keeping the other parameters as listed in Ref. 35 (see Supporting Material for more details).

## RESULTS

### Strain stiffening can be observed for several Ndc80 variants

The ability of different Ndc80 variants to track and stall depolymerizing MTs has been examined in two previous studies (7, 10). Here, we re-examined the underlying data to determine effective stiffnesses for wild type Ndc80 complexes (Ndc80^wt^) as well as three variants, including Ndc80 complexes with a phosphorylated tail (Ndc80^P^), Ndc80 complexes, whose tail has been truncated (Ndc80^Δ80^), and Ndc80 complexes with mutated CH domains and a greatly reduced MT binding (Ndc80^CHmut^).

In Ref. 7, the stiffnesses for the wild type Ndc80 were determined from the variance of bead positions during MT stall, which gave one stiffness per experiment. These experiments revealed stiffening under force. However, the stall force is related to the Ndc80 density on the bead and therefore probably also to the number of Ndc80 complexes that are attached to the MT. For this reason, we can not determine whether the stiffening is an intrinsic property of the Ndc80 complex and/or the PF, or whether it arises from a higher number of parallelly attached Ndc80 complexes.

To settle this issue, we need to time-trace the stiffness throughout a *single* experiment. We achieve this by binning the data of bead positions in intervals of 0.1 s and determining an effective stiffness and applied force for each interval separately using the procedure described in the Materials and Methods section. This allows us to determine the stiffness while the depolymerizing MT pulls the attached bead and thereby increases the force. In other words, we measure the stiffness for different forces in a single experiment and, thus, with a fixed Ndc80 density on the bead. The results of this analysis are shown in Figs. 3A-D for four representative examples, as well as in Fig. S1 in the Supporting Material for twelve further examples.

**Figure 3:**
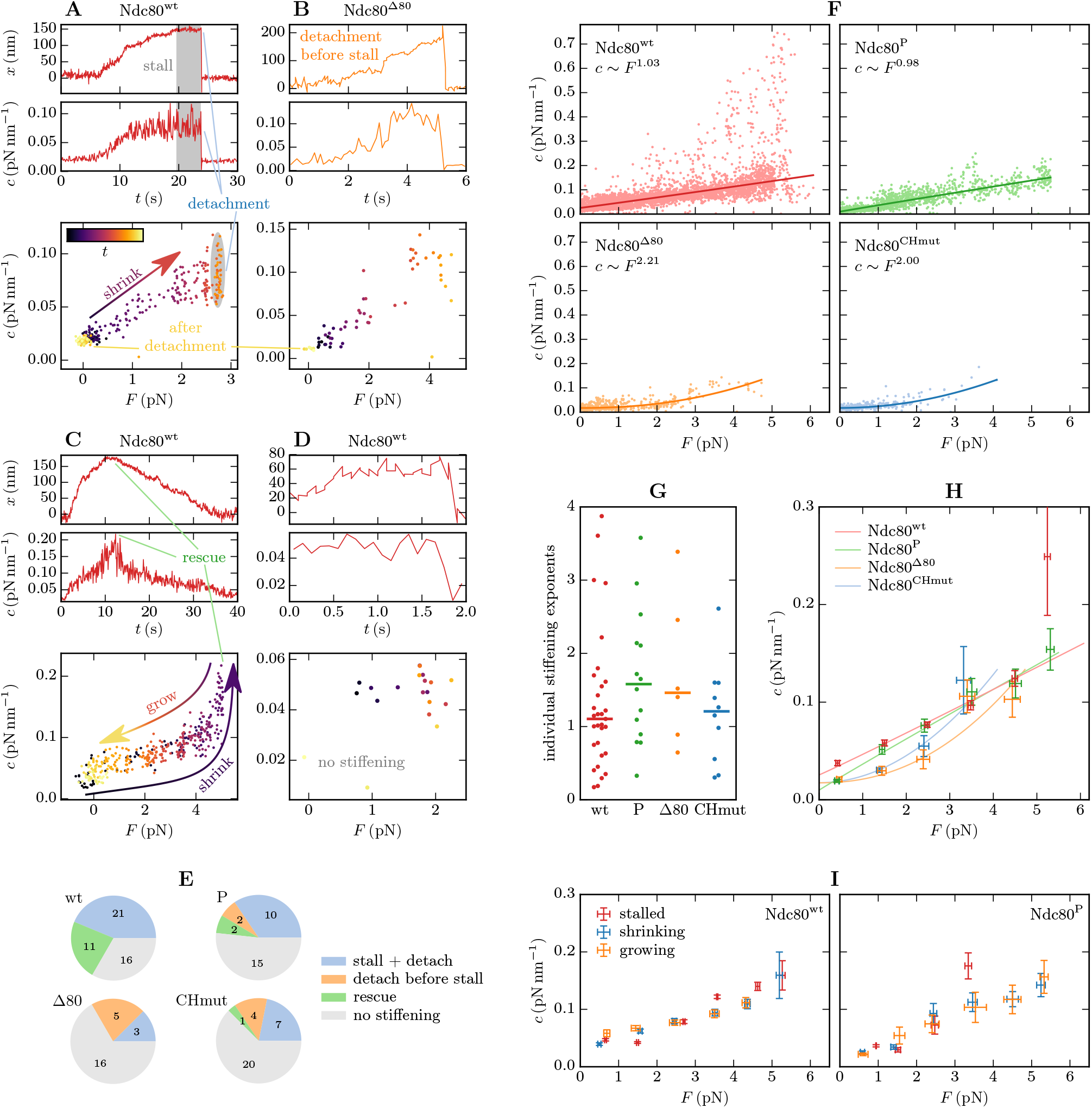
(A-D) Four representatives for typical behavior during single experiments. Further examples, including Ndc80^CHmut^ and Ndc80^P^, are shown in Fig. S1 in the Supporting Material. For each experiment, the upper two plots show the piecewise linearized bead position and the stiffness over time, while the stiffness–force relation is depicted in the bottom plot, in which the time is represented by the color gradient. When the force is increased due to the shrinking MT, we can observe stiffening in most experiments (A-C). Shrinking usually is followed by a stall phase with constant force and constant stiffness (gray), except for experiments with Ndc80^Δ80^ (B) and Ndc80^CHmut^, in which the bead often detaches before reaching the stall force. The stall is terminated either by detachment (A) or by MT rescue (C). After detachment the bead relaxes to *x* = 0 and we measure the trap stiffness. When the MT is rescued and grows, the stiffness–force relation traces the same curve as during depolymerization. (E) Observed frequencies of the typical events described in (A-D). (F) Robust power fits. For each Ndc80 variant, the combined stiffness data from all experiments (dots) is robustly fitted with power functions *c*(*F*) = *aF*^*m*^ + *c*_0_ (lines) as described in the Supporting Material. The offset *c*_0_ should respect the trap stiffness. (G) Stiffening exponents *m* that result from applying the power fits to those single experiments, where a proper stiffening is recognizable and the bead was attached to the MT for at least 1 s, so that there are ≥ 10 stiffnesses to fit to. The horizontal bars mark the medians. (H) Summarized stiffness–force relations. The error bars represent the standard deviation of the mean in *F* and in *c* of the points within each bin, where each experiment is considered as an independent event. The lines show the power fits from (F). (I) Stiffnesses separately evaluated for stalled, shrinking, and growing MTs. Error bars as described in (H). Ndc80^Δ80^ and Ndc80^CHmut^ are not included here because of rare occurrence of rescues.

For each of the Ndc80 variants, there are experiments in which we detect a smooth stiffening behavior when the tension increases, but there are also experiments that do not exhibit any stiffening at all as in Fig. 3D. Based on previously observed absence of stiffening during lateral bead–MT attachment (7), we interpreted experiments without stiffening as laterally attached and excluded them from the following analyses. We note that our main results and conclusions that we derive in the following are unchanged if the experiments without stiffening are included, see Fig S2 in the Supporting Material. Experiments with observed stiffening were interpreted as resulting from the end-on attachment that is sketched in Fig. 2A and analyzed further. Stiffening was observed in 32 of 48 experiments with Ndc80^wt^, 8/24 with Ndc80^Δ80^, 12/32 with Ndc80^CHmut^, and 14/29 with Ndc80^P^ (see Fig. 3E). Calibrations of QPD voltage to obtain bead displacement, as well as calibrations of trap stiffness as a function of bead’s displacement from the trap center were linear within 200 nm from the trap center. Therefore, displacements of a bead from the trap center that exceeded 200 nm were excluded from the following analyses of the strain stiffening.

In the experiments that exhibit stiffening, the MT depolymerization typically stalls at some force for Ndc80^wt^ and Ndc80^P^, but not for Ndc80^Δ80^ and Ndc80^CHmut^, where the bead often detaches before it can stall the MT, see Fig. 3B, and the stalls that occur are shorter (10). This indicates that the detachment force for Ndc80^Δ80^ and Ndc80^CHmut^ is usually below the MT stall force under the experimental conditions applied. Therefore, a systematic analysis of these mutants is only enabled by time-tracing of the stiffness prior to detachment, as measurements in a stalled state are rarely possible. The decrease in detachment force also confirms that the combination of the tail and the CH domain is essential for force-resisting attachment (10).

A stall can be interrupted in two possible ways: Either the bead detaches from the MT (Fig. 3A), or MT growth is rescued (Fig. 3C). In the first case, the bead relaxes immediately to the equilibrium position of the optical trap and the measured stiffness is the trap stiffness *c*_trap_. In case of a rescue, the bead stays attached to the polymerizing MT and the tension decreases until the MT undergoes a catastrophe and starts depolymerizing again. This will allow us to investigate stiffening of the Ndc80 complexes in dependence of the dynamic state of the MT in the following section. The four typical behaviors (stiffening during stall followed by detachment or rescue, detachment before stall, or no stiffening) that are described above and in Figs. 3A-D can be observed for each of the Ndc80 variants with frequencies as summarized in Fig. 3E.

Another advantage of our time-tracing method is that we generate many more data points from the same number of experiments. While, for example, 53 stiffnesses of Ndc80^wt^ were determined in Ref. 7 from stalling MTs, time-tracing generates 4403 stiffnesses from the same experiments (see Fig. S3 in the Supporting Material). The cumulated force–stiffness data, which are collectively depicted in Fig. 3F for each Ndc80 variant, reveal the strain stiffening more clearly than the sole use of the stalls in Ref. 7 and allow for a more thorough analysis and interpretation.

In order to further characterize strain stiffening, we fit power law functions *c*(*F*) = *aF*^*m*^ + *c*_0_ to the collective data by use of a robust regression minimizing the Huber loss (36), see Fig. 3F and Supporting Material. The exponent *m* characterizes the observed strain stiffening behavior. We account for the trap stiffness by the offset *c*_0_ for which the fits yield results that, indeed, lie in a range 0.01 pN nm^*-*1^ to 0.025 pN nm^*-*1^, which is in accordance with the trap stiffnesses determined during calibration. While the stiffnesses exhibit a linear dependence on force with exponents *m* around unity for Ndc80^wt^ and Ndc80^P^, the stiffness–force relations of Ndc80^Δ80^ and Ndc80^CHmut^ have a roughly parabolic shape (*m* ≈ 2). We note that for Ndc80^Δ80^ and Ndc80^CHmut^, there are few data for high forces, making the stiffening exponents less reliable.

The stiffening exponents determined in Fig. 3F are the result of a fit to the combined data from several experiments, and may be biased by the correlation between the stall force and the Ndc80 densities on the bead. Our new time-tracing analysis, however, dampens this bias compared to the sole analysis of the stalls as stiffnesses are determined for small forces but at high Ndc80 densities during MT depolymerization in the respective experiments. We also apply the power law regressions to the stiffness–force relations of single experiments. Then, we find most stiffening exponents in a range between 0.5 and 2, but also detect outliers below this range and close to 4, Fig. 3G. Finally, we evaluated the stiffness–force relations separately for different populations of beads. Since beads within the same population were prepared under the same conditions, they can be assumed to be coated with a similar number of Ndc80 complexes. Though the populations had very different mean Ndc80 densities, the separate evaluation does not reveal a correlation between Ndc80 density and the stiffening behavior as shown by Fig. S4 in the Supporting Material. Therefore, we exclude that the linear stiffening is a result of the correlation between stall force and number of Ndc80 complexes per bead.

For a better visibility and comparability of the strain stiffening of the different Ndc80 variants, we summarize the stiffness measurements in Fig. 3H by binning the time-traced stiffnesses of all experiments in bins of 1 pN width and averaging force and stiffness in each of these bins. We find that all variant Ndc80 complexes exhibit similar strain stiffening. Both truncating the tail or modifying the CH domain slightly decrease the stiffness compared to the wild type, especially for small forces. This indicates that in all four Ndc80 variants, the remaining intact common parts play a central role in strain stiffening. Differences between variants are due to changes in the Ndc80–MT bond and will be addressed in the Discussion section.

### Measured stiffness is independent of MT state

When a MT is rescued after the stall, as in the experiment depicted in Fig. 3C, the time-traced stiffness follows the same stiffness–force relation in the shrinking and growing states before and after rescue, respectively. The same observation can be made in experiments with several consecutive rescues and catastrophes. This indicates that the stiffness is independent of the MT state, i.e., on whether the Ndc80 complexes are attached to a shrinking, stalled, or growing MT. To address this question systematically, we identify the phases of shrinking, stalled and growing MTs in the single experiments, and bin the stiffnesses analogously to Fig. 3H but separately for each of the three MT states. The results in Fig. 3I do not show a significant dependence on the MT state for any of the four Ndc80 variants. The independence of Ndc80-MT linker stiffness from MT state indicates that PFs either do not contribute to the total stiffness (because *c*_PF_ ≫ *c*_tot_), or that, if they do contribute, their mechanical properties do not depend on whether the MT is growing or shrinking. The stiffness contributions of PFs are investigated below in more detail.

### PLL-PEG is too stiff to contribute to the measured stiffnesses

In the following sections, we attempt to explain the observed strain stiffening by modeling the elastic elements between the bead and the MT that add up to the stiffness 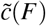 as shown in Fig. 2B and Eq. 2. We start with the PLL-PEG linkage to the bead, whose contribution to the measured stiffnesses should be clarified in addition to those from the Ndc80 complex or the PF. PLL-PEG consists of a poly-L-lysine (PLL) backbone and polyethylene glycol (PEG) side chains. While the PLL backbone is adsorbed on the glass bead in the experiment, the flexible PEG chains can bind the Ndc80 oligomers.

From force–extension curves measured in AFM experiments (37), stiffness of PLL adsorbed on a flat Nb_2_O_5_ surface can be estimated as ∼100 pN nm^*-*1^ which is three orders of magnitude above the stiffnesses observed in our experiments. Oesterhelt *et al*. (35) successfully described the elasticity of single PEG polymers by means of an extended FJC model with a bond length that follows from a two-level system. Applying this model to the PEG-2000 with *N*_S_ = 45 segments used in our experiments, we obtain a force-free stiffness of 1.38 pN nm^*-*1^, that increases under tension as shown in the Supporting Material.

We conclude that both the PLL backbone and the PEG chains are too stiff to make a significant contribution to the observed strain stiffening and can be omitted in the following analysis (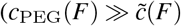 in Eq. 2.

### Stiffening is a direct consequence of the Ndc80 structure

After eliminating a possible contribution of the PLL-PEG, we will next include the remaining elements (Ndc80 complex, MT, PF) step by step to discuss their relevance for the observed strain stiffening. We start with the Ndc80 complex, which is the most promising and interesting candidate because of its flexible structure and its force transmitting role in the mitotic spindle.

We model the Ndc80 complex as a FJC with two flexibly connected stiff arms (see Fig. 2C), resulting in a stiffness *c*_Ndc_(*F*) as given by Eq. 11. We have to take into account that there might be *n* Ndc80 complexes attached in parallel. For this situation, we assume that each Ndc80 complex has the same elongation so that the force *F* is shared equally by the attached complexes and the total stiffness is given by *nc*_Ndc_(*F/n*) (see Eq. 3). In Fig. 4A, we compare the total stiffness of *n* parallel Ndc80 complexes to the measured stiffnesses of Ndc80^wt^. For any *n*, the model results are about a factor 2-4 larger than the measured values. Moreover, the two-segment Ndc80 model results in a strain stiffening exponent *m* = 2 in Eq. 12, which is larger than the measured values *m* ≈ 1 in Fig. 3E. We conclude that the stiffness of the Ndc80 complex alone can explain strain stiffening but is not sufficient to explain the measured stiffnesses quantitatively.

**Figure 4:**
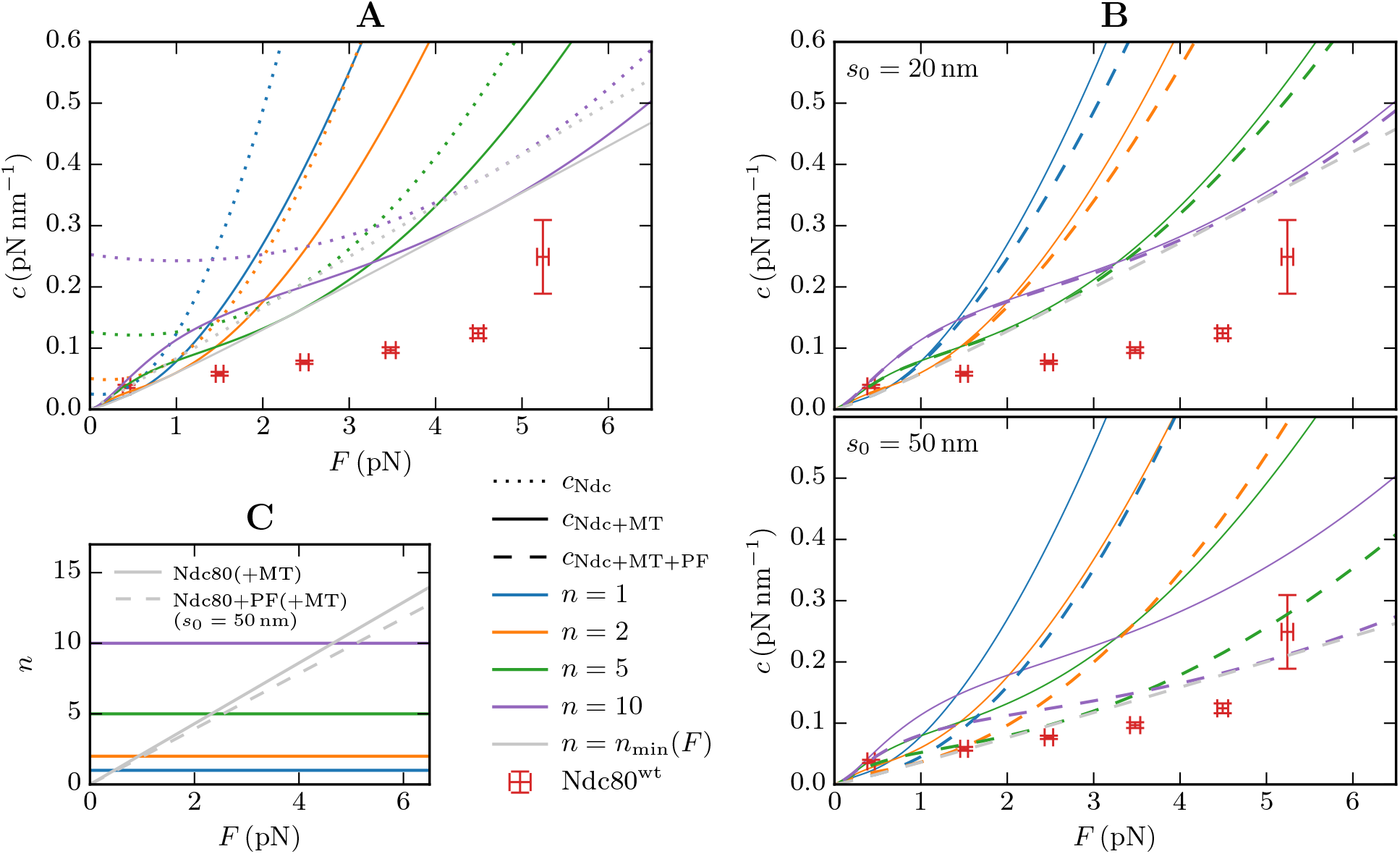
(A) The stiffness of *n* parallel Ndc80 complexes *nc*_Ndc_(*F/n*) according to Eq. 10 (dotted lines) is by a factor of 2–4 larger than the measurements with wild type Ndc80. Adding the MT stiffness in series (solid lines) reduces the stiffness, but not sufficiently to match the measurements. The theoretical stiffness is minimized when the number of attached Ndc80 complexes increases linearly with the force (*n*_min_(*F*), gray line). (B) Stiffness of a MT in series with *n* parallel Ndc80 complexes, each of which is in series with a PF (dashed lines). Top: When the attachment on the curved PF has a distance of *s*_0_ = 20 nm from the straight MT lattice, the stiffness only differs slightly from the stiffness of a MT in series with just the Ndc80 complexes (solid lines). Bottom: For *s*_0_ = 50 nm the PF does have a significant contribution to the total stiffness. However, the combined stiffnesses of Ndc80 complexes and PFs are still larger than the experimental outcomes for wild type Ndc80. The Ndc80^wt^ values in (A) and (B) are not corrected by the trap stiffness (see text). (C) Force-dependent number of attached Ndc80 complexes that minimizes the theoretical stiffness (*n*_min_(*F*), gray).

We note that the experimental values in Fig. 4 still contain the trap stiffness, which is not included in the theoretical curves. We refrained from correcting the experimental values for the trap stiffness, as trap stiffness and time-traced stiffnesses were determined by different methods and are, therefore, difficult to compare: while the time-traced data was obtained from the equipartition theorem as described in the Materials and Methods section, the trap stiffness, which was also used for force determination, was calibrated from the power spectral density (PSD) (38–40). In conclusion, one should keep in mind that the wild type stiffness values in Fig. 4 probably need to be shifted slightly downwards by an amount of approximately 0.02 pNnm^*-*1^.

The fact that the Ndc80 model gives larger stiffnesses than measured motivates the consideration of the additional elastic elements from the MT, the PFs and the Ndc80–MT bonds in the total stiffness in Eq. 4. These elements are in series to the Ndc80 complex and potentially reduce the total stiffness of the entire Ndc80–MT link.

### Entropic MT stiffness

Next, we consider the contribution of the MT, which can be attached to several Ndc80 complexes. While a purely mechanical MT stiffness that follows from stretching or bending the MT can be ruled out to contribute significantly, see Supporting Material, the MT also has an entropic elasticity from thermal fluctuations. To estimate the entropic stiffness, we describe the MT as a worm-like chain by means of the force–extension relation in Eq. 5 taking into account a length-dependent persistence length (29). The relevant MT length is the free part of the MT, i.e., the contour length between the fixed MT seed and the MT tip. As a further consequence of the length-dependent persistence length, the resulting MT stiffness in Eq. 6 becomes length-independent for small MT lengths *L* ≪ *L*_0_ = 21 µm. Therefore, the MT stiffness can be assumed to be the same in all experiments, where typical MT lengths are between 3 and 10 µm, and we use *L* = 10 µm in the following as a general representative.

The entropic stiffness of the MT is of the same order of magnitude as the Ndc80 stiffness, which is somewhat surprising for a MT that is by a factor 100 shorter than its persistence length. Consequently, the stiffness of *n* parallel Ndc80 complexes alone is significantly reduced when the MT stiffness is added in series, see Fig. 4A. Interestingly, the combination of a single MT with stiffening exponent 3/2 and several (*n* ≳, 5) Ndc80 complexes stiffening with *F*^2^ results in a roughly linear behavior in the examined force range, which might explain the measured stiffening exponents in Fig. 3E. Despite the overall stiffness reduction, our model predictions are still too large as compared to the measurements when only the MT and the Ndc80 complexes are considered. Therefore, we speculate in the next step that the Ndc80 complexes bind to the curved parts of the PFs resulting in an effective stiffness from straightening the PFs.

### Only long protofilaments are flexible enough to reduce the total stiffness

We assume that forces recorded in our experiments are produced by tubulin protofilaments bending outwards from the MT lattice when a MT shortens. We always record stall events after a force has been developed, i.e. we do not observe bead-bound Ndc80 preventing MT shortening without force development. Therefore, we hypothesize that bead-bound Ndc80 interacts with bent protofilaments during stall. We also assume that during stall, the MT-generated force is equalized by the returning force of the optical trap directed along the MT axis in the opposite direction.

Since the PFs are bent out of their preferred curvature, a restoring force on the Ndc80 complexes builds up, governed by an effective stiffness *c*_PF_ for strains in the axial direction. In the Materials and Methods section, we quantified this stiffness with a simple beam model (see Fig. 2D) in Eq. 14. This model depends on three parameters: the preferred curvature *ϕ* of the PF, its persistence length *L*_p_ that defines the bending stiffness *α* = *k*_B_*TL*_p_, and the position *s*_0_ along the free part of the PF where the Ndc80 complex attaches, i.e., where the force is applied. While the first two parameters as well as the length of the free curved parts of the PF are well described for MTs in vitro (34), a fixed attachment position *s*_0_ of Ndc80 complex to the PF is not known, and we use typical values *s*_0_ = 20 nm and *s*_0_ = 50 nm.

In Fig. 4B, we plotted the stiffnesses of *n* parallel Ndc80 complexes with *c*_Ndc_ that are each in series with a PF *c*_PF_. We see that the influence of PF bending is negligible for *s*_0_ = 20 nm, whereas for *s*_0_ = 50 nm the PF significantly reduces the total stiffness close to the measured wild type values. We conclude that the PF stiffness is only relevant, when the Ndc80 complex is attached near the end of a sufficiently long PF. An upper bound for *s*_0_ is given by the total length of the curved part of the PF for which values of *L* = 10-80 nm have been measured in vitro (34).

### Catch bond behavior lowers the stiffness of parallel Ndc80 complexes

So far, we have assumed a *fixed* number *n* of parallel Ndc80 complexes over the whole range of applied forces. We now analyze how our model behaves when the number *n* of attached Ndc80 complexes depends on the applied force, *n* = *n*(*F*). In principle, both a catch bond and a slip bond mechanism are conceivable for the Ndc80–MT bond. A catch (slip) bond is characterized by a binding affinity that increases (decreases) under tension (41). In our model, a catch (slip) bond implies a monotonically increasing (decreasing) force-dependent number of attached Ndc80 complexes *n*(*F*).

We find that a force-dependent *n*(*F*) can lower the effective total stiffness of the entire Ndc80–MT link. The minimal cooperative stiffness that can be realized is the envelope of the stiffness–force relations *c*_*n*_(*F*) for different *n* in Fig. 4AB. The envelope of the *c*_*n*_(*F*) relations is obtained by a force-dependent number *n*_min_(*F*) of attached Ndc80 complexes that minimizes the stiffness *c*_*n*_(*F*) for each *F*. The resulting *n*_min_(*F*) is shown in Fig. 4C and increases linearly, which also implies a catch bond behavior. For long PFs (*s*_0_ = 50 nm), the envelope stiffness *c*_*n*_min (*F*) has actually the correct order of magnitude as compared to the experimental data shown in Fig. 4B. The number *n*_min_(*F*) also remains below *n* = 15 in the examined force range, which is in agreement with estimates for the number of Ndc80 complexes in the proximity of the MT given the expected Ndc80 density on the bead (5, 7).

Apart from the absolute stiffness values, a model should reproduce the observed roughly linear strain stiffening behavior with the exponent *m* around unity (see Fig. 3E). Our theory, on the other hand, predicts exponents of 3/2, 2 and 5/2 for the MT, the Ndc80 complexes and the PFs, respectively. The relation *n*(*F*) can change the latter two stiffening exponents. From Eq. 3, one can see that a linearly increasing *n*(*F*) actually implies a linear stiffening as observed in our experiments since *F/n*(*F*) = const and, therefore, 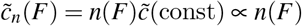.

For a slip bond, on the other hand, where *∂*_*F*_ *n*(*F*) < 0 and *n*(*F*) → 0 for large forces by definition, the argument in Eq. 3, *F/n*(*F*), increases rapidly. As a consequence, we can approximate *c*_Ndc_ and *c*_PF_ with the power laws from Eqs. 12 and 15, respectively, so that

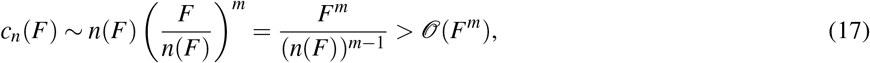

We conclude that any kind of slip bond will increase the stiffening exponent for large forces. Stiffening exponents below the values *m* = 2 for Ndc80 stiffness or *m* = 2.5 for PF stiffness, as obtained in the experiments, are indicative of a catch bond mechanism. For the time traces from single experiments in Fig. 3F, 79 % of the stiffening exponents lie below 2.

## DISCUSSION

The novel analysis of time traces of bead positions in optical trapping experiments allows us to time-trace the stiffness during single experiments and go beyond an analysis limited to the MT stall state (7). This enabled us to increase the available number of stiffness measurements by 1 to 2 orders of magnitude and study strain stiffening of wild type Ndc80 in more detail. It also enabled us to study strain stiffening of Ndc80 variants with a truncated tail or with a mutated CH domain, both of which typically detach before the MT has been stalled, and to explore strain stiffening selectively in the shrinking, stalled and growing state of MTs.

We found that all mutant Ndc80 complexes exhibit strain stiffening hinting at a central role of the Ndc80–Nuf2 coiled coil, rather than the MT binding sites in the CH-domain and the N-terminal tail. The measured stiffnesses of the entire Ndc80–MT link are lower than theoretical predictions from our Ndc80 model. Therefore, additional elastic elements in the Ndc80–MT link or a force-dependent number of attached Ndc80 complexes should be relevant. We could exclude the PLL-PEG connection between Ndc80 complexes and the bead, which is too stiff to contribute. Effects from the MT and from protofilament flexibility are necessary for the model to conform to experimentally measured stiffnesses, and the observed linear stiffening may be explained by an increasing number of attached Ndc80 complexes under force (catch bond). Moreover, stiffness differences between variants allow us to draw some conclusions on the Ndc80–MT bond.

### Effects from protofilaments

Time-tracing of the stiffness allowed us to study strain stiffening selectively in the shrinking, stalled and growing state of MTs. We found no significant stiffness differences between MT states (see Figs. 3C and H) leading to the conclusion that the stiffness of the bent PFs is either too large to contribute significantly to the total stiffness or that it does not depend on the MT’s polymerization state.

Our PF model showed that the PF contribution to the overall stiffness depends strongly on the length of the PFs and the position along the PF where the Ndc80 complex is attached. An attachment point *s*_0_ close to the straight part of the PF (*s*_0_ ≤ 20 nm) predicts a negligible contribution of the PF. It has been shown that single Ndc80 complexes bind to straight PFs but show a much weaker affinity for curved PFs (6, 31). The results of Ref. 31 also suggest that the weaker affinity is due to a missing interaction of the CH domain with a curved PF so that a weaker bond is formed by the N-terminal tail alone. This is supported by the results in Fig. 3E, which show that Ndc80^Δ80^ with truncated tail enters a state of stall, which requires binding to the MT, less frequently. There are, however, differences. Our stiffness measures derive from experiments where Ndc80 multimers were used that were able to tip-track—in contrast to the single Ndc80 complexes in Refs. 6, 31. Moreover, these experiments were not performed with full wild type Ndc80 complexes but with Ndc80^bonsai^, lacking almost the entire coiled coil region (31), or with Ndc80^broccoli^, lacking Spc24 and Spc25 (6). Together, these differences leave room for speculating that our experimental conditions allow for Ndc80 binding to curved PFs. For instance, the Ndc80 complexes may bind initially to a straight PF, which becomes curved at the binding site during depolymerization while the Ndc80 complex stays attached.

Attachment of Ndc80 to long flaring PFs with attachment lengths around *s*_0_ = 50 nm are predicted to reduce the stiffness of the Ndc80–MT link to values close to the measured stiffnesses (Fig. 4B). This is consistent with the conclusion that PFs contribute to the total stiffness but have identical elastic properties during MT shrinkage, stall and growth, and is also in agreement with the recent observation that the curvature of PFs is the same during polymerization and depolymerization (34).

### Number of attached Ndc80 complexes and potential catch bond mechanism

The number *n* of simultaneously attached Ndc80 complexes could not be directly inferred from the experiments in Refs. 7, 10. From the known Ndc80 densities on the bead we can estimate that 1 to 4 Ndc80 trimers, i.e., 3 to 12 Ndc80 complexes are in the vicinity of the MT end and can potentially bind (7).

From the equal stiffness–force relations before and after a rescue, we conclude that also the number of attached Ndc80 complexes is the same during polymerization and depolymerization. Dynamically, it is possible that the number of attached Ndc80 complexes changes in a force-dependent manner between MT catastrophes and rescues. For force-free detachment and attachment of a single Ndc80^wt^ complex, time scales of *τ*_off_ = 1.6 s and *τ*_on_ = 0.4 s, respectively, have been found (7). With a phosphorylated tail, *τ*_off_ is supposed to be smaller (12). Since the durations of MT depolymerization vary in a wide range from less than 1 s up to ∼100 s, both a constant and a dynamic number of attached complexes are possible.

Our modeling results showed that the absolute stiffness values as well as the roughly linear strain stiffening relation (see Fig. 3E) are best reproduced by a force-dependent number *n*(*F*), which increases linearly with force (see Fig. 4B) implying a catch bond mechanism. Among the stiffening exponents for the time traces from single experiments in Fig. 3F, 79 % of the strain stiffening exponents lie below 2, which is indicative of a catch bond like behavior. As already noted above, the linear stiffening may also be an apparent phenomenon that results in the examined force range from the combination of the different stiffening behaviors of a single MT and a sufficiently large constant number of Ndc80 complexes and PFs.

Assuming that Ndc80 forms a catch bond to the MT, the question arises how this mechanism could work. The whole kinetochore, consisting of several additional proteins, has been proposed to act like a catch bond (42). It is widely assumed that the Aurora B kinase, which was not present in our experiments, is important in the kinetochore’s catch bond mechanism (43–45). Kinetochores purified from S. cerevisiae, however, were shown to build catch bonds with MTs in vitro even without Aurora B activity (46). Importantly, the catch-bond behavior of the purified yeast kinetochores was abolished when Dam1 or Stu2 were absent, even in presence of Ndc80 (46, 47). There is evidence that Ndc80 stretching correlates with MT binding as the Ndc80 complex may exist in an auto-inhibited bent conformation with reduced MT binding capacity (48) and that the Ndc80 complex bends (“jackknifes”) upon detachment (49). This suggests that the binding affinity for MTs increases when the Ndc80 complex is stretched, which would further imply that Ndc80 has an intrinsic catch-bond-like mechanism. To what extent this effect is relevant for our experiments is unclear, as according to our FJC model the Ndc80 complex is already stretched by small forces. For instance, at *F* = 1 pN we find a mean angle of (44 ± 23)° between the arms *a* and *b* (where the angle is 0° for a maximally stretched Ndc80 complex). Speculating on other possible catch bond mechanisms, it is conceivable that by stretching the unstructured tail additional binding sites become available which are concealed in the entangled tail at low forces. Force-enhanced adhesion by unfolding is a common catch bond mechanism (50); examples are the von Willebrand factor (51, 52) and *α*-catenin bonding to F-actin (53) in the cytoskeleton. In conclusion, an intrinsic catch bond mechanism of Ndc80 complexes is compatible with current knowledge and worth considering as one of the ingredients that define the characteristics of Ndc80 strain stiffening. Such a mechanism could also contribute to the catch bond behavior of the entire kinetochore.

We assumed throughout the paper that the applied force is shared equally among the Ndc80 complexes attached in parallel so that each complex and each PF has the same elongation. If this assumption is lifted, the system will be dominated by a few very stiff Ndc80 complexes, which are the ones with the largest stretch. It is thinkable that the linker extensions approach a uniform distribution under force, for instance because some extremely stretched Ndc80 complexes detach from the shrinking MT. Then, despite detachment of a few linkers, the number of linkers with a relevant stretch increases under force. In conclusion, such a mechanism would result in an apparent catch bond behavior without the need for individual Ndc80–MT catch bonds.

### Structure of Ndc80–MT bond and role of the N-terminal tail in MT binding

Stiffness differences between the wild type Ndc80^wt^ and mutants Ndc80^Δ80^ and Ndc80^CHmut^ reflect roles in MT binding of the the N-terminal tail and the CH domain, respectively. These differences will allow us to draw some conclusions on the Ndc80–MT bond, whose exact mechanism is still unknown. The CH domain binds between two tubulin monomers via a structural element named the “toe”(31). The N-terminal tail also supports MT binding, but its exact role is still under debate. Three models are currently discussed (23, 54): direct binding to the MT lattice, cooperativity and clustering by interactions with neighboring Ndc80 complexes, and co-factor recruitment. Since no co-factors were present in our experiments, we will concentrate on the former two models.

Our comparative study of Ndc80^wt^, Ndc80^Δ80^ and Ndc80^CHmut^ allows us to dissect the role of the tail and the CH domain on Ndc80–MT binding. Both Ndc80 mutants displayed decreased stiffnesses compared to the wild type (Fig. 3G), an effect we ascribed either to a softened Ndc80–MT bond or to a reduced number of attached Ndc80 complexes. In the elastic toy model in Fig. 5A, both the CH domain and the tail of Ndc80^wt^ bind to the PF, so that they can be represented by two parallel springs that add up to the total bond stiffness *c*_bond_. Due to the parallel arrangement, lack of either of the two springs in Ndc80^Δ80^ or Ndc80^CHmut^ reduces the bond stiffness, predicting overall stiffnesses as depicted in Fig. 3G. If both the CH domain and the N-terminal tail bind directly to the MT lattice, the two Ndc80 mutants probably have a reduced MT binding affinity. Therefore, in the direct binding model, the reduction of the overall stiffness could be a consequence of a reduced number of attached Ndc80 complexes as well as of a reduction of the individual bond stiffnesses.

**Figure 5:**
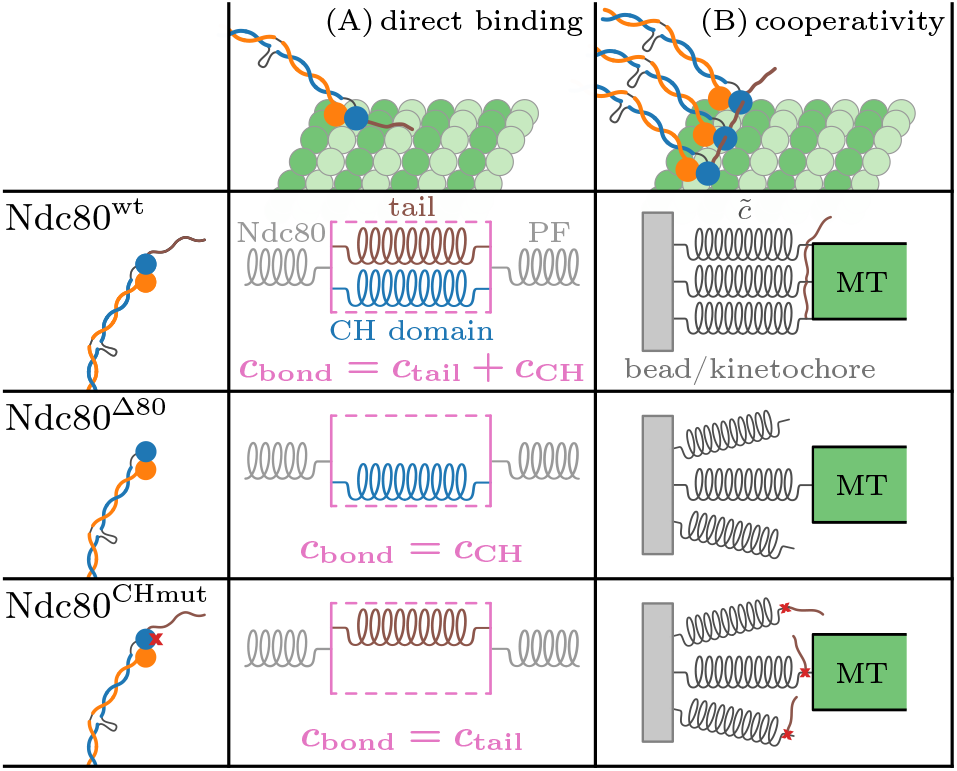
Models of the Ndc80–MT bond for potential roles of the N-terminal tail. (A) When the tail binds directly to the PF, an elastic model of the Ndc80–MT bond comprises the tail and the CH domain as two parallel springs. If the tail is deleted (Ndc80^Δ80^) or the CH binding domain is blocked (Ndc80^CHmut^), either of those springs is missing which lowers the bond stiffness *c*_bond_ and thereby the total stiffness 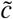. (B) In the case of an interaction of the tail with CH domains of neighboring Ndc80 complexes, the number of parallelly attached linkers is reduced for the two mutants Ndc80^Δ80^ and Ndc80^CHmut^.

When the N-terminal tail interacts with neighboring Ndc80 complexes instead of the tubulin, the induced cooperativity supports parallel attachments of multiple Ndc80^wt^ complexes, see Fig. 5B. Deletion of the tail or mutation of the CH domain may impair the cooperative behavior, with the result that fewer linkers connect the bead (or the kinetochore) with the MT and the overall stiffness decreases (for small and medium forces, see Fig. 4). While there is evidence supporting a contribution of the tail to cooperative MT binding, there is at present no clear evidence indicating that the Ndc80 CH domain contributes to binding cooperativity.

Tail phosphorylation was shown to reduce the MT binding affinity of the Ndc80 complex (2, 11–14). Moreover, the durations of the MT stalls are shorter when the Ndc80 tails are phosphorylated (10). This suggests that phosphorylation impedes the function of the tail similarly as if the tail is truncated. However, in contrast to Ndc80^Δ80^, the stiffness of Ndc80^P^ is not significantly reduced compared to Ndc80^wt^, see Fig. 3G. It remains an open question how these observations are compatible with the binding models depicted in Fig. 5.

### Comparison of model and experiment

Our model can explain the strain stiffening from the structure of the Ndc80 complex and the bending elasticity of long flaring PFs. If we allow for a force-dependent number of bound Ndc80 complexes, a linearly increasing number of bound complexes (requiring a catch bond mechanism) also reproduces the observed roughly linear strain stiffening relation. Ndc80, by itself or in series with PFs, gives absolute stiffness values that are still above the measured stiffnesses. Including a force-dependent number of bound Ndc80 complexes gives values which are close but still slightly higher than measured. As already mentioned, the missing part might be the Ndc80–MT bond, which is additionally in series with the Ndc80 complex and the PF as illustrated in Fig. 2. However, since the stiffness of the bond is hard to quantify due to the little knowledge of the exact binding mechanism, we did not go beyond the qualitative discussion in the previous section.

Regarding the Ndc80, we have also investigated possible effects from finite flexibility of the two Ndc80 arms by modeling both arms as semiflexible worm-like chains. We performed Monte Carlo simulations, in which each Ndc80 arm had the same persistence length *L*_p_. Using persistence lengths in a realistic range of *L*_p_ ≳, 100 nm as they have been determined for other coiled coil proteins such as tropomyosin (55), our simulations lead to the conclusion that semiflexibility has only a minor effect on total stiffness (see Supporting Material).

Finally, we need to discuss the accuracy of our experimental method. While the trap stiffness was calibrated by fitting the power spectrum based on Brownian motion in a harmonic potential (38–40), this well-established method can not be applied for measurements of the anharmonic Ndc80 stiffness. Therefore, we had to determine the Ndc80 stiffness from the variance of bead position, which is a simpler and universal but less accurate approach. In particular, the measured overall variance contains a systematic noise that is added to the actual variance of bead position, which is why the stiffness tends to be underestimated by *k*_B_*T/*Var(*x*) (56–58). This systematic underestimation, which we can not quantify, might explain why our model tends to predict higher stiffnesses than experimentally measured.

## CONCLUSION

In conclusion, we accumulated extensive stiffness data of the Ndc80–MT link by optical trapping methods in combination with a novel time-tracing analysis. We were able to study wild type Ndc80 complexes and three variants. Our theoretical model for the Ndc80–MT link, which includes structure-based models of the Ndc80 complex, the MT and flaring PFs, is able to explain the strain stiffening observed when Ndc80 is bound to shortening MT plus ends, and reproduces the correct order of magnitude of the stiffness. Thus, our results on the elastic properties further support these structural models from the mechanical point of view. Our model also reproduces the roughly linear strain stiffening behavior when taking a force-dependent binding affinity into account.

## AUTHOR CONTRIBUTIONS

A.M., M.D., V.A.V. and P.J.H. conceived and designed the experiments. P.J.H. contributed reagents. V.A.V. performed and analyzed the experiments. J.K. and F.S. developed the theoretical model. F.S. analyzed the data and performed calculations. F.S. and J.K. wrote the original draft. A.M., V.A.V. and P.J.H. reviewed and edited the manuscript.

## DECLARATION OF INTEREST

The authors declare no competing interests.

## ACKNOWLEDGMENTS

V.A.V. and M.D. acknowledge support from European Research Council Synergy Grant MODELCELL (proposal 609822). A.M. acknowledges funding by European Research Council (ERC) through Synergy Grant 951439 (BIOMECANET).

## SUPPORTING MATERIAL

Supporting Material can be found online at http://www.biophysj.org.

## SUPPORTING CITATIONS

References 59-61 appear in the Supporting material.

## Notes

### Competing Interest Statement

The authors have declared no competing interest.

### Summary of Updates

sections Theoretical model and Results updated to include MT stiffness

