## Supporting Material for "Strain stiffening of Ndc80 complexes attached to microtubule plus ends"

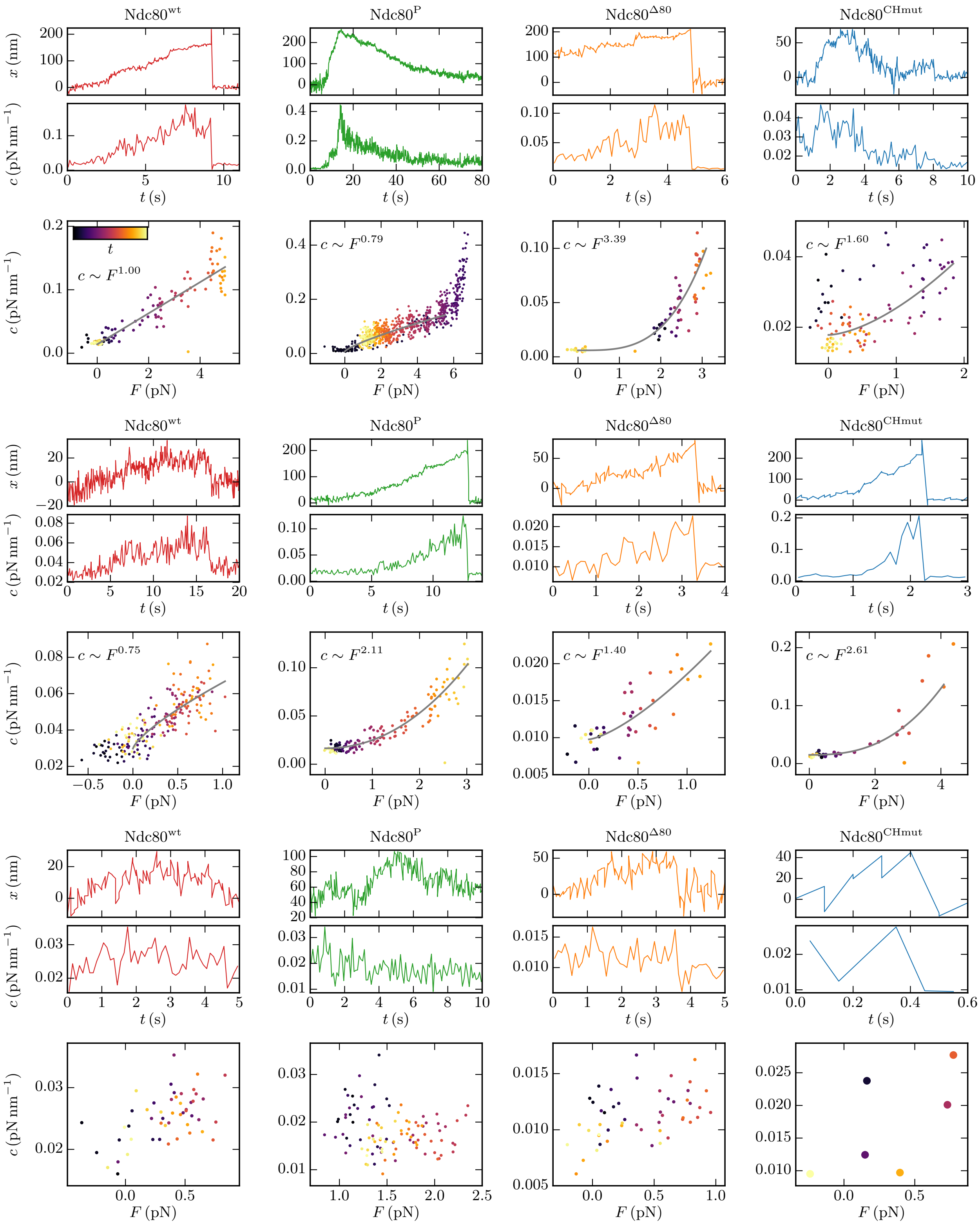


FIG. S1. Twelve more example experiments as described in Fig. 3A-D in the main text. The gray lines show powerfits to the stiffness–force relations that exhibit strain stiffening, giving the stiffening exponents depicted in Fig. 3F in the main text. The fits do not include values corresponding to $x > 200\,\mathrm{nm}$, see for instance the top Ndc80$^{\mathrm{P}}$ plot, where the fit only applies to $F \lesssim 6\,\mathrm{pN}$.

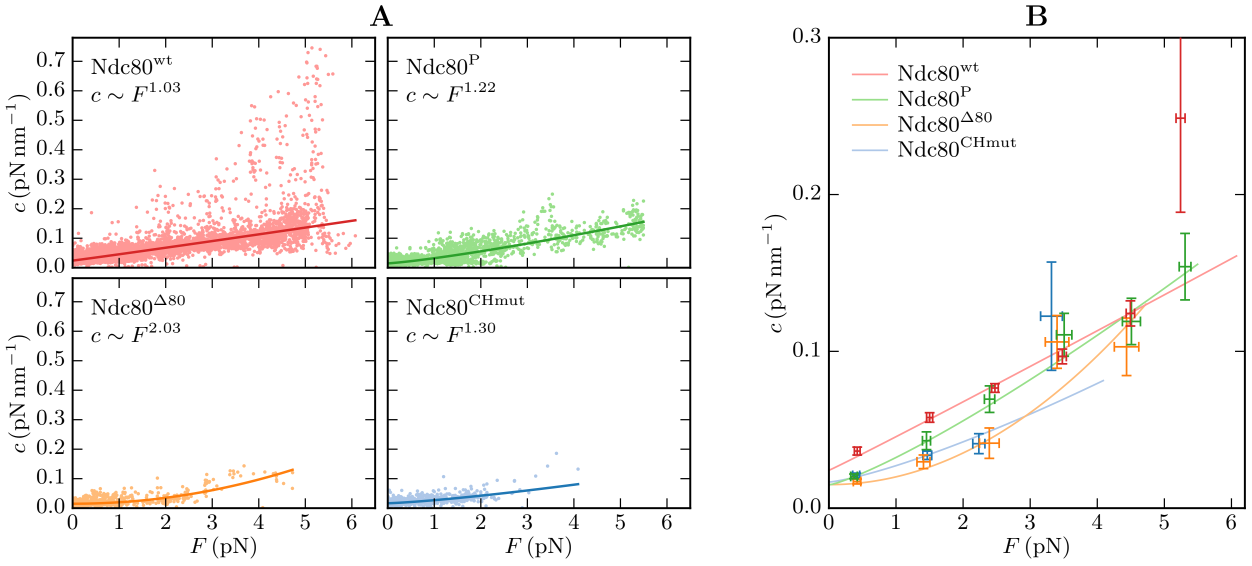


FIG. S2. Collective analysis including the experiments without observable stiffening. The main results are unchanged due to the comparably short durations of non-stiffening experiments. (A) Cumulative stiffness data with robust power fits analogously to Fig. 3F in the main text. The linear stiffening is maintained for Ndc80$^{wt}$ and roughly for Ndc80$^{P}$. (B) Summarized stiffness–force relations analogously to Fig. 3G in the main text. The Ndc80$^{wt}$ stiffness still exceeds the stiffnesses of Ndc80$^{\Delta 80}$ and Ndc80$^{CHmut}$.

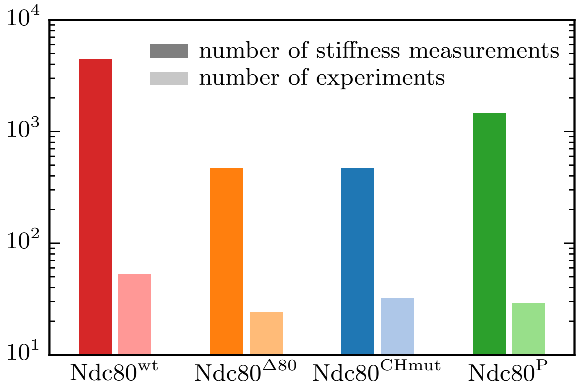


FIG. S3. Increased amount of stiffness data. While only one stiffness per experiment can be obtained by the sole use of the stalls for stiffness determination (right bars), the time tracing analysis increases the amount of data by 1 to 2 orders of magnitude (left bars).

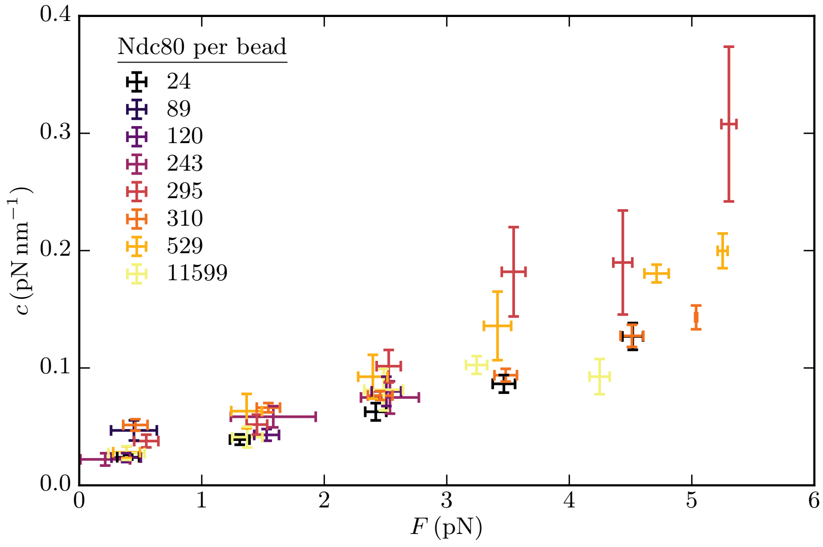


FIG. S4. Stiffness–force relations for different bead populations. The legend shows the mean number of Ndc80 complexes per bead in each population. There is no recognizable correlation between the stiffness and the Ndc80 density. The binning was performed as in Fig. 3H in the main text.

### I. ROBUST FITTING TO THE STIFFNESS–FORCE RELATIONS

Here, we describe the robust fitting procedure of power law functions $c(F) = aF^b + c_0$ to the force–stiffness data ($F_i$,$c_i$). Since the stiffness values that result from the time tracing analysis exhibit a high spread (see Fig. 3E in the main text), we perform the power fits by use of a robust fitting method instead of a least squares fit which is sensitive to outliers. While in a least squares fit the sum of the squared residuals is minimized,

$$\min \frac{1}{2} \sum_i \epsilon_i^2, \quad \epsilon_i = \frac{c(F_i) - c_i}{\sigma_i}, \tag{S1}$$

the robust fitting method that we applied minimizes the sum of the Huber loss functions $\rho(\epsilon_i)$ [S1]:

$$\min \sum_i \rho(\epsilon_i), \quad \rho(\epsilon) = \begin{cases} \frac{1}{2}\epsilon^2, & |\epsilon| < k \\ k|\epsilon| - \frac{1}{2}k^2, & |\epsilon| \geq k \end{cases}. \tag{S2}$$

Thereby, the residuals are weighted as in a least squares fit for $\epsilon < k$, and with an absolute estimator (which results in a median when the data is fitted by a constant function) for $\epsilon \geq k$. The tuning parameter $k$ is set to 1.345 to achieve a relative efficiency of 95 % in respect to the normal distribution [S2, S3]. Finally, we need to estimate the errors $\sigma_i$ of the stiffnesses that we determined from the sample variances of the bead positions $x$ within each interval (see Fig. 1 in the main text). Since $c \propto 1/\mathrm{Var}(x)$, the relative deviation of the the stiffness is the same as for the variance, $\sigma_c/c = \sigma_{\mathrm{Var}}/\mathrm{Var}(x)$. From a sample of length $n$, the variance and its deviation can be estimated to [S4]

$$\mathrm{Var}(x) = \frac{1}{n-1} \sum_i \left(x_i - \langle x \rangle\right)^2, \quad \sigma_{\mathrm{Var}} = \sqrt{\frac{2}{n-1}}\,\mathrm{Var}(x). \tag{S3}$$

With $n = 1000$ in a 0.1 s interval at a sample frequency of 10 kHz and stiffnesses around $c = 0.1\,\mathrm{pN\,nm^{-1}}$, the estimated error is

$$\sigma_c = \sqrt{\frac{2}{n-1}}\,c = 0.0045\,\mathrm{pN\,nm^{-1}}. \tag{S4}$$

The fit results and the original data are shown in Fig. 3E in the main text, the fit parameters are listed in Tab. S1.

TABLE S1. Fit parameters of power fits $c(F) = aF^b + c_0$.

| | $a$ | $b$ | $c_0$ |
|---|---|---|---|
| $\mathrm{Ndc80}^{\mathrm{wt}}$ | 0.022 | 1.03 | 0.025 |
| $\mathrm{Ndc80}^{\Delta 80}$ | 0.0038 | 2.21 | 0.017 |
| $\mathrm{Ndc80}^{\mathrm{CHmut}}$ | 0.0069 | 2.00 | 0.017 |
| $\mathrm{Ndc80}^{\mathrm{P}}$ | 0.027 | 0.98 | 0.010 |

### II. MECHANICAL MT STIFFNESS

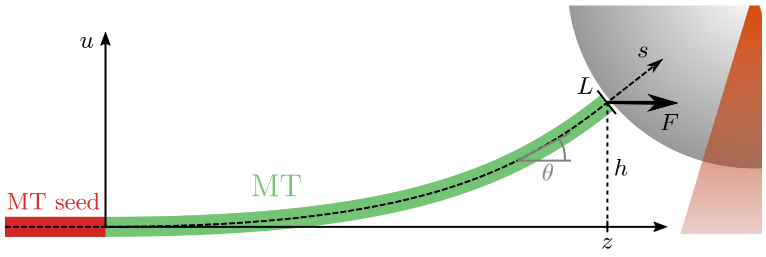


FIG. S5. Model for the effective stiffness from MT unbending.

The model for the mechanical stiffness that follows from MT unbending is similar to the PF model in the main text. In contrast to the PF, the MT does not have a spontaneous curvature but its tip is lifted to a height $h$ by the attached bead as sketched in Fig. S5. With a force $F$ that is applied in $z$-direction, the total energy is given by the sum of bending and stretching energies:

$$E_{\mathrm{MT}} = \int_0^L \left(\frac{\alpha}{2}(\dot{\theta}(s))^2 - F\cos\theta(s)\right) \mathrm{d}s, \tag{S5}$$

where $s$ denotes the position along the MT, $\theta$ the local bending angle, $L$ the length of the free MT end (behind the fixed MT seed), and $\alpha = k_{\mathrm{B}}TL_{\mathrm{p}}$ the bending stiffness. The configuration $\theta(s)$ that minimizes the energy under the constraint

$$h = \int_0^L \sin\theta(s)\,\mathrm{d}s \tag{S6}$$

satisfies the Euler-Lagrange equation

$$\ddot{\theta} = \frac{F}{\alpha}\sin\theta - \frac{F_h}{\alpha}\cos\theta \overset{\theta \ll 1}{\approx} \frac{F}{\alpha}\theta - \frac{F_h}{\alpha}, \tag{S7}$$

with a Largange-multiplier $F_h$, which corresponds to the force in $u$-direction that is necessary to hold the MT tip at height $h$. With the boundary conditions $\theta(0) = 0$ (due to the fixed MT seed) and $\dot{\theta}(L) = 0$, the approximated equation is solved by

$$\theta(s) = \frac{F_h}{F}\left(1 - \cosh(\lambda s) + \tanh(\lambda L)\sinh(\lambda s)\right), \quad \lambda := \sqrt{\frac{F}{\alpha}}, \tag{S8}$$

$$F_h = \frac{Fh/L}{1 - \frac{1}{\lambda L}\tanh(\lambda L)}. \tag{S9}$$

This results in the effective deflection

$$\begin{aligned} z &= \int_0^L \cos\theta(s)\,\mathrm{d}s \approx \int_0^L \left(1 - \frac{\theta^2(s)}{2}\right)\mathrm{d}s \\ &= L\left[1 - \frac{h^2}{4L^2}\left(\frac{3}{1 - f(\lambda L)} - \left(\frac{\lambda L\, f(\lambda L)}{1 - f(\lambda L)}\right)^2\right)\right], \quad f(x) := \frac{\tanh x}{x}. \end{aligned} \tag{S10}$$

Finally, the effective stiffness of MT unbending is given by

$$
\begin{aligned}
c &= \left(\frac{\partial z}{\partial F}\right)^{-1} \\
&= \frac{8\sqrt{F\alpha}}{h^2} \frac{\left(1 - f(\lambda L)\right)^3}{f'(\lambda L)\left(\left(3 + 2\lambda^2 L^2\right) f(\lambda L) - 3\right) + 2\lambda L f^2(\lambda L)\left(1 - f(\lambda L)\right)}.
\end{aligned}
\tag{S11}
$$

In the limit of strong forces ($F \gg \alpha/L^2 \sim 0.05\,\mathrm{pN}$), this converges to

$$
c \approx \frac{8L^2}{h^2\sqrt{\alpha}} F^{\frac{3}{2}}. \tag{S12}
$$

There are two required parameters: the length $L$ of the free MT end and the height $h$ of the MT tip. Typical MT lengths lay around $L \sim 5\,\mathrm{\mu m}$. Given the bead diameter of $1\,\mathrm{\mu m}$ and the distance between the surfaces of the coverslip and the bead ($\sim 100\,\mathrm{nm}$), the height of the tip should not exceed $h \approx 500\,\mathrm{nm}$. Fig. S6 shows the mechanical MT stiffness $c_{\mathrm{MT}}$ compared to and in series with the stiffness of $n$ parallel Ndc80 complexes for $L = 5\,\mathrm{\mu m}$ and $h = 500\,\mathrm{nm}$. Though this choice of $h$ provides a lower estimate ($c_{\mathrm{MT}} \propto h^{-2}$), the mechanical MT stiffness is still too large, to have a significant influence.

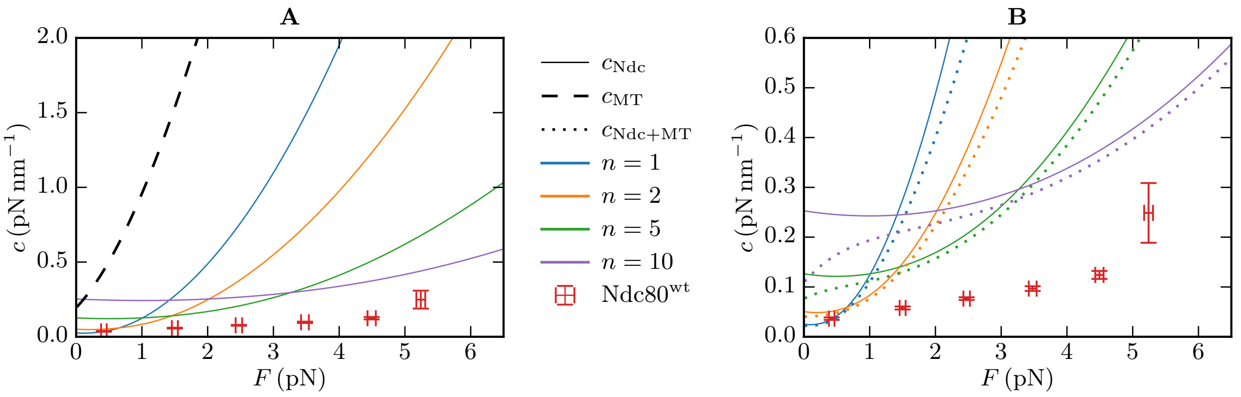


FIG. S6. (A) The mechanical stiffness of MT unbending (dashed line, $L = 5\,\mathrm{\mu m}$, $h = 500\,\mathrm{nm}$) exceeds the stiffness of $n$ parallel Ndc80 complexes (solid lines) and the measured wild type stiffnesses. (B) The dotted lines show the mechanical MT stiffness in series with $n$ Ndc80 complexes. The influence of the MT is not significant when $c_{\mathrm{Ndc}} \gg c_{\mathrm{MT}}$.

### III. PEG STIFFNESS

According to Oesterhelt et al. [S5], the extension–force relation of PEG is given by:

$$L(F) = N_{\mathrm{S}} \left( \frac{L_{\mathrm{planar}}}{\mathrm{e}^{+\beta\Delta G(F)} + 1} + \frac{L_{\mathrm{helical}}}{\mathrm{e}^{-\beta\Delta G(F)} + 1} \right) \left( \coth(\beta F L_{\mathrm{K}}) - \frac{1}{\beta F L_{\mathrm{K}}} \right) + N_{\mathrm{S}} \frac{F}{K_{\mathrm{S}}}, \qquad \text{(S13)}$$

where $\Delta G(F) = \Delta G_0 - F\,(L_{\mathrm{planar}} - L_{\mathrm{helical}})$. The used parameters are listed in Tab. S2. Except $N_{\mathrm{S}}$, they are taken from Ref. [S5]. Fig. S7 shows the stiffness $c_{\mathrm{PEG}}(F) = (\partial L(F)/\partial F)^{-1}$ compared to and in series with the Ndc80 stiffness.

TABLE S2. Parameters of the PEG model.

| Description | Symbol | Value |
|---|---|---|
| number of segments | $N_{\mathrm{S}}$ | 45 |
| bond length of the planar (trans-trans-trans) conformation | $L_{\mathrm{planar}}$ | 0.358 nm |
| bond length of the helical (trans-trans-gauche) conformation | $L_{\mathrm{helical}}$ | 0.28 nm |
| Kuhn length | $L_{\mathrm{K}}$ | 0.7 nm |
| zero force free energy difference $G_{\mathrm{planar}} - G_{\mathrm{helical}}$ | $\Delta G_0$ | $3\,k_{\mathrm{B}}T$ |
| segment elasticity | $K_{\mathrm{S}}$ | $1.5 \times 10^5\,\mathrm{pN\,nm^{-1}}$ |

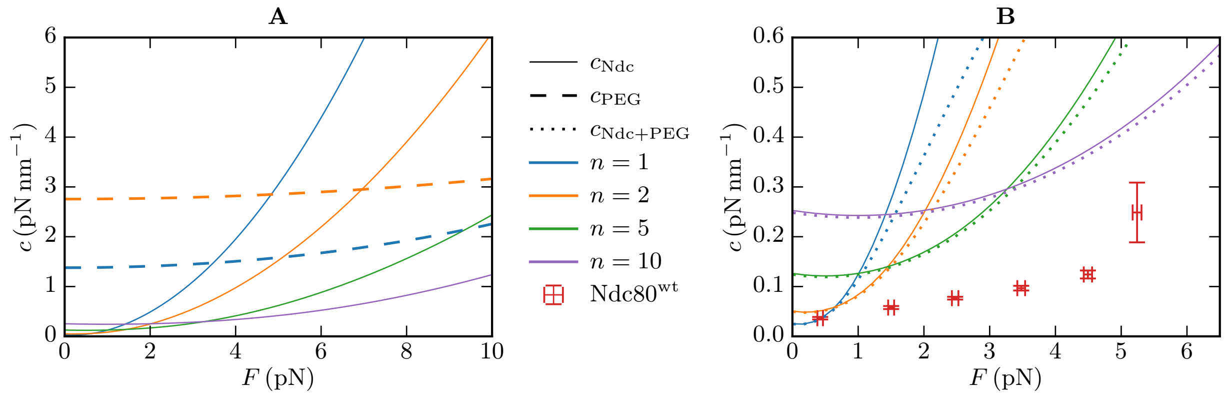


FIG. S7. (A) The stiffness of $n$ parallel PEG molecules (dashed lines) exceeds the Ndc80 stiffness (solid lines) in most of the examined force range ($F < 6$ pN) and lies an order of magnitude above the measured stiffnesses ($\sim 0.1\,\mathrm{pN\,nm^{-1}}$). (B) The dotted lines show the stiffness of $n$ PEG molecules in series with $n$ Ndc80 complexes. The influence of PEG is not significant when $c_{\mathrm{Ndc}} \gg c_{\mathrm{PEG}}$.

### IV. WORM-LIKE CHAIN MODEL

To take into account a Ndc80 complex with possibly (semi)flexible arms, we model the Ndc80 complex as two worm like chains (WLC) with persistence length $L_\mathrm{p}$ that are flexibly connected. To run Monte Carlo (MC) simulations, we describe each WLC by a bead–spring model [S6]: The Ndc80 arm $\vec{a}$ ($\vec{b}$) is discretized into $N_a + 1$ ($N_b + 1$) beads, which are connected by $N_a$ ($N_b$) springs each with rest length $d_a$ ($d_b$) and stiffness $k_a$ ($k_b$). The rest lengths are defined by $d_a = a/N_a$ and $d_b = b/N_b$ to be consistent with the observed Ndc80 arm lengths $a$ and $b$. Given the extensions and directions of the springs as $\vec{a}_i$ ($\vec{b}_j$) with $i = 1...N_a$ ($j = 1...N_b$), we find the total stretching energy

$$E_\mathrm{stretch} = \frac{k_a}{2}\sum_{i=1}^{N_a}\left(|\vec{a}_i| - d_a\right)^2 + \frac{k_b}{2}\sum_{j=1}^{N_b}\left(|\vec{b}_j| - d_b\right)^2. \tag{S14}$$

The positions $\vec{A}_m$ and $\vec{B}_n$ of the beads are given by

$$\vec{A}_m = \vec{A}_0 + \sum_{i=1}^{m}\vec{a}_i, \qquad \vec{B}_n = \vec{B}_0 + \sum_{j=1}^{n}\vec{b}_j, \tag{S15}$$

where $\vec{A}_0 = \vec{0}$ (the Ndc80 complex is fixed to the glass bead) and $\vec{B}_0 = \vec{A}_{N_a}$. The glass bead, which is modeled as a wall (see Fig. 2C in the main text), is described by the boundary condition that each bead has to be located in the upper half space, i.e., for the $z$-components: $A_{m,z}, B_{n,z} > 0$ for each $m, n$. Each bead $\vec{A}_m, \vec{B}_n$ except $\vec{A}_0$, $\vec{A}_{N_a} = \vec{B}_0$ and $\vec{B}_{N_b}$ contributes a bending energy which sums up to

$$E_\mathrm{bend} = \frac{L_\mathrm{p}}{\beta d_a}\sum_{m=1}^{N_a-1}(1-\cos\alpha_m) + \frac{L_\mathrm{p}}{\beta d_b}\sum_{n=1}^{N_b-1}(1-\cos\beta_n), \tag{S16}$$

where $\alpha_m$ ($\beta_n$) is the angle between bonds $m$ and $m+1$ ($n$ and $n+1$):

$$\cos\alpha_m = \frac{\vec{a}_m\cdot\vec{a}_{m+1}}{|\vec{a}_m||\vec{a}_{m+1}|}, \quad \cos\beta_n = \frac{\vec{b}_n\cdot\vec{b}_{n+1}}{|\vec{b}_n||\vec{b}_{n+1}|}. \tag{S17}$$

Finally, when a force $F$ in $z$-direction is applied on the last bead, the total energy reads as:

$$E = E_\mathrm{stretch} + E_\mathrm{bend} - FB_{N_b,z}. \tag{S18}$$

In the MC simulation, we randomly choose a bead ($i, j > 0$) and suggest to move it in a random direction for a constant distance $s$. If the move does not violate the boundary condition, it will be accepted with probability $\min(1, \exp(-\beta\Delta E))$, where $\Delta E = E_\mathrm{sugg} - E_\mathrm{orig}$

is the energy difference between the suggested and the original configuration following Eq. (S18). After a certain time of equilibration, we measure $z = B_{N_b,z}$ and $z^2$ after each sweep ($N_a + N_b$ moves), and calculate the mean extension and its variance at the end of a simulation. By repeating it for various external forces, we can record force–extension and force–stiffness relations.

In our simulations, we used $k_a = k_b = 1000\,\mathrm{pN\,nm^{-1}}$ to model the two Ndc80 arms as nearly unstretchable. Moreover, we used the same discretization lengths $d_a = d_b = 1\,\mathrm{nm}$, i.e., $N_a = 40$ and $N_b = 16$.

Fig. S8 shows the stiffness–force relations for a single and for $n = 5$ parallel semiflexible Ndc80 complexes with various persistence lengths $L_\mathrm{p}$. The effect of semiflexibility on the stiffness is negligible for realistic persistence lengths above 100 nm [S7].

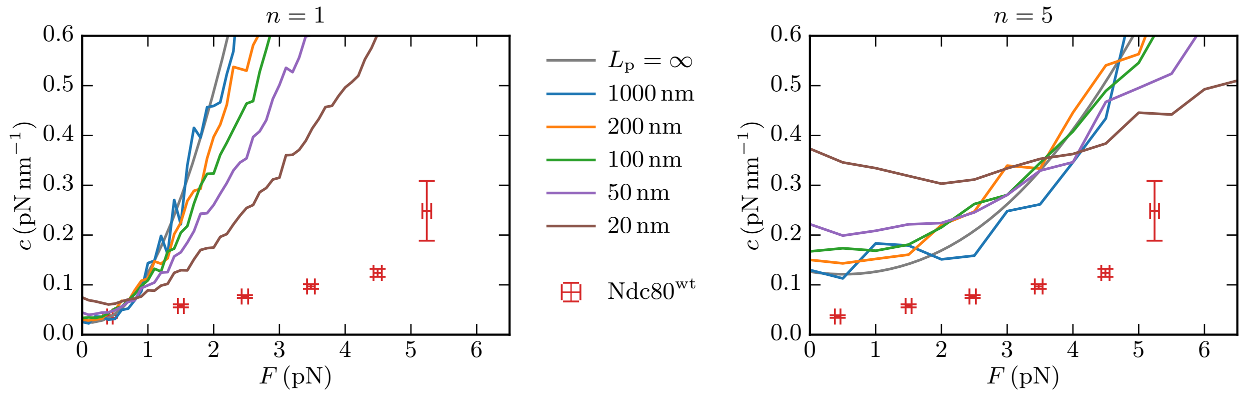


FIG. S8. Stiffness of a single (left) and $n = 5$ parallel (right) semiflexible Ndc80 complexes with persistence lengths $L_\mathrm{p}$. As a check of the MC simulations, we added the large persistence length $L_\mathrm{p} = 1000\,\mathrm{nm}$, which correctly resembles results of the FJC model ($L_\mathrm{p} = \infty$) according to Eq. (8) in the main text.

[S4] Casella, G., and R. L. Berger, 2002. Statistical inference. Duxbury, Pacific Grove, CA 93950 USA, 2 edition.

[S5] Oesterhelt, F., M. Rief, and H. E. Gaub, 1999. Single molecule force spectroscopy by AFM indicates helical structure of poly(ethylene-glycol) in water. *New Journal of Physics* 1:6.1–6.11. https://iopscience.iop.org/article/10.1088/1367-2630/1/1/006.

[S6] Kierfeld, J., O. Niamploy, V. Sa-yakanit, and R. Lipowsky, 2004. Stretching of semiflexible polymers with elastic bonds. *The European Physical Journal E* 14:17–34. http://link.springer.com/10.1140/epje/i2003-10089-3.

[S7] Hvidt, S., J. D. Ferry, D. L. Roelke, and M. L. Greaser, 1983. Flexibility of light meromyosin and other coiled-coil $\alpha$-helical proteins. *Macromolecules* 16:740–745. https://pubs.acs.org/doi/abs/10.1021/ma00239a007.
